# HiMAP: robust Phylogenomics from Highly Multiplexed Amplicon sequencing

**DOI:** 10.1101/213454

**Authors:** Julian R. Dupuis, Forest T. Bremer, Angela Kauwe, Michael San Jose, Luc Leblanc, Daniel Rubinoff, Scott M. Geib

## Abstract

High-throughput sequencing has fundamentally changed how molecular phylogenetic datasets are assembled, and phylogenomic datasets commonly contain 50-100-fold more loci than those generated using traditional Sanger-based approaches. Here, we demonstrate a new approach for building phylogenomic datasets using single tube, highly multiplexed amplicon sequencing, which we name HiMAP (Highly Multiplexed Amplicon-based Phylogenomics), and present bioinformatic pipelines for locus selection based on genomic and transcriptomic data resources and post-sequencing consensus calling and alignment. This method is inexpensive and amenable to sequencing a large number (hundreds) of taxa simultaneously, requires minimal hands-on time at the bench (<1/2 day), and data analysis can be accomplished without the need for read mapping or assembly. We demonstrate this approach by sequencing 878 amplicons in single reactions for 82 species of tephritid fruit flies across seven genera (384 individuals), including some of the most economically-important agricultural insect pests. The resulting dataset (>150,000 bp concatenated alignment) contained >40,000 phylogenetically informative characters, and although some discordance was observed between analyses, it provided unparalleled resolution of many phylogenetic relationships in this group. Most notably, we found high support for the generic status of *Zeugodacus* and the sister relationship between *Dacus* and *Zeugodacus*. We discuss HiMAP, with regard to its molecular and bioinformatic strengths, and the insight the resulting dataset provides into relationships of this diverse insect group.

## INTRODUCTION

High throughput sequencing has transformed the status quo methodologies for many biological disciplines, and phylogenomics is no exception (Lemmon and Lemmon 2013, McCormack, *et al.* 2013). Although whole-genome phylogenies are still out of reach for most non-model systems, targeted and reduced representation sequencing approaches (“genomic partitioning”) can generate datasets that are orders of magnitude larger than “traditional” Sanger sequencing-based molecular phylogenetic datasets. The sheer size of these targeted genomic datasets provides unprecedented resolution into phylogenetic relationships (Blaimer, *et al.* 2015, Leache, *et al.* 2015, Prum, *et al.* 2015, Dupuis, *et al.* 2017), particularly when the methodological and analytical difficulties of such large datasets are considered (Philippe, *et al.* 2011, Kumar, *et al.* 2012, Xi, *et al.* 2015). The various approaches for genomic partitioning have their own strengths and weaknesses (reviewed in McCormack, *et al.* (2013) and Lemmon and Lemmon (2013)), including cost, phylogenetic depth (species-level or deeper relationships), the number of individuals to be sequenced, analytical limitations, and resulting data type and locus length (single nucleotide polymorphism (SNP) vs. sequence data). For example, restriction-site associated DNA sequencing (RAD-seq) provides a cost-effective approach for sequencing hundreds of individuals, but is limited to relatively recent scales of divergence (due to restriction enzyme site variation), and SNP-based or short sequence-based (generally <100 bp) datasets that can limit appropriate analyses (Leache, *et al.* 2015, DaCosta and Sorenson 2016, Dupuis, *et al.* 2017). On the other hand, transcriptome-based approaches can generate large datasets of long sequence-based loci, but become expensive with more than a few dozen samples and require high quality RNA input, which is often limiting (Hedin, *et al.* 2012, Johnson, *et al.* 2013, Kawahara and Breinholt 2014).

For generating sequence-based phylogenomic datasets with long loci, sequence capture approaches (also called “target enrichment” or “hybrid enrichment”) are at the forefront of the field. Variations of sequence capture, such as anchored hybrid enrichment (AHE: Lemmon, *et al.* 2012) and the use of ultra-conserved elements (UCEs: Faircloth, *et al.* 2012b), use a set of modified oligo probes to capture genomic DNA of interest. Particularly in the early stages of these methods, probe sets were quite expensive; as a way to make these approaches more economically feasible, probe sets were often developed for order-level or higher groups of organisms and shared by multiple experiments (e.g. Hymenoptera (Blaimer, *et al.* 2015, Faircloth, *et al.* 2015, Blaimer, *et al.* 2016a), Amniota (Faircloth, *et al.* 2012b, Ruane and Austin 2017), Vertebrata (Lemmon, *et al.* 2012, Brandley, *et al.* 2015, Peloso, *et al.* 2016)), and these higher-level probe sets continue to be used for both deep- and shallow-scale phylogenomics. Library preparation for these methods involve several main steps including shearing/fragmentation, sequencing library construction (adapter addition), enrichment/sequence capture, and final pooling and quantification, and generally take multiple days of bench time. Most studies have targeted <100 specimens (e.g. Lemmon, *et al.* 2012, McCormack, *et al.* 2012, Faircloth, *et al.* 2013, Hedtke, *et al.* 2013, Blaimer, *et al.* 2016a, Hamilton, *et al.* 2016, Hosner, *et al.* 2016, Breinholt, *et al.* 2017), however, datasets sampling 100-200 specimens have also been generated (e.g. Prum, *et al.* 2015, Moyle, *et al.* 2016, Branstetter, *et al.* 2017a, Branstetter, *et al.* 2017b).

Here we present a novel and cost-effective approach for generating phylogenomic datasets of hundreds to thousands of genes and hundreds of individuals using amplicon sequencing based on highly multiplexed polymerase chain reaction (PCR). Multiplex PCR simultaneously targets multiple loci by including multiple primer pairs in a single reaction (Chamberlain, *et al.* 1988), but its use for developing high throughput sequencing libraries has been hindered by many challenges. Primary among these are the difficulty in amplifying more than a few dozen targets, time-intensive optimization of reaction conditions, uneven and off-target amplification, and formation of primer dimers (Edwards and Gibbs 1994, Markoulatos, *et al.* 2002, Fan, *et al.* 2006, Turner, *et al.* 2009). In a phylogenomic context, these challenges can be exacerbated by sequence variation at priming sites and target length variation (Lemmon and Lemmon 2013), thus limiting phylogenetic applications of multiplex PCR to relatively shallow time scales and few targets (e.g. Phuc, *et al.* 2003, Stiller, *et al.* 2009, Doumith, *et al.* 2012, Wielstra, *et al.* 2014). Amplicon sequencing has been used to generate moderate sized phylogenomic datasets, but has generally relied on singleplex, barcoded PCR products being pooled into high throughput sequencing libraries (O'Neill, *et al.* 2013, Barrow, *et al.* 2014), or microfluidic systems that facilitate automated amplification of singleplex or multiplex reactions (Richardson, *et al.* 2012, Gostel, *et al.* 2015, Uribe-Convers, *et al.* 2016).

We adapt a new library preparation procedure originally developed for human cancer research, and demonstrate its first use in a phylogenomic context by sequencing 878 conserved exons for 384 specimens of tephritid fruit flies (Diptera: Tephritidae). Tephritidae includes some of the most economically important pest species in the world (White and Elson-Harris 1992, Vargas, *et al.* 2015); despite large research efforts for control and management of these pests, many morphologically-cryptic species complexes remain uninvestigated and there is a general lack of consensus regarding main relationships between genera (Hendrichs, *et al.* 2015, Virgilio, *et al.* 2015, Schutze, *et al.* 2016). We focus our specimen sampling on genera across Tephritidae that contain some of the most economically important pests, including *Anastrepha*, *Bactrocera*, *Ceratitis*, and *Zeugodacus*, as well as on the morphologically-cryptic complexes within *Bactrocera* and *Zeugodacus*. Our 878-exon panel uses a single oligonucleotide primer pool for all of these genera, and we demonstrate that this approach remedies virtually all of the main difficulties in using multiplex PCR for high-throughput sequencing library construction (Turner, *et al.* 2009, Lemmon and Lemmon 2013), particularly for shallow-mid scale phylogenies. This approach is cost-effective, and library preparation can be accomplished in <1/2 day. The approach developed here for data processing to call consensus sequences is rapid and straightforward, avoids read mapping and assembly, and can be accomplished using a basic laptop or desktop computer. We name this approach HiMAP (Highly Multiplexed Amplicon-based Phylogenomics).

## METHODS

### Overview of end-to-end HiMAP approach

We present an end-to-end concept for completing a HiMAP project, including methods for locus selection, primer design, target amplification, sequencing, and post-sequencing data processing and analysis (henceforth we use “data processing” to refer to the steps going from raw sequence data to input files for phylogenetic analyses). The locus selection and data processing pipelines are summarized in Figure 1 and Figure 2, respectively. We demonstrate this with a focus on a moderate phylogenetic time scale, using genera in Tephritidae diverging 65-100 million years ago (Krosch, *et al.* 2012, Caravas and Friedrich 2013), but a deeper or more shallow focus could be taken, depending on the research question. We use thirteen genomic and transcriptomic data sources for locus selection, spanning three genera. However, the pipeline presented here is amenable to fewer or more data sources, as long as one of them has relatively high quality genome assembly and structural annotation of protein coding regions. Additionally, loci could be selected with other methods and integrated into the later HiMAP steps. Wet lab requirements are standard, using common equipment found in most molecular labs (thermocycler, fluorometer, magnetic stand). Sequencing depth needed to generate a high-quality consensus for several hundred loci across hundreds of species can be reached with low-cost bench-top sequencing (e.g. Illumina MiSeq), and computational requirements for post-sequencing data processing (going from raw FASTQ files to aligned consensus sequences) are minimal, and could be completed using a laptop or desktop computer. Finally, the output formatting is standard multi-FASTA format, so can be readily used or easily re-formatted for routine phylogenetic analyses. Details of bioinformatic pipelines (including locus selection, amplicon/primer filtering, and post-sequencing data processing) and additional code used here can be found at https://github.com/popphylotools/HiMAP.

**Figure 1.**
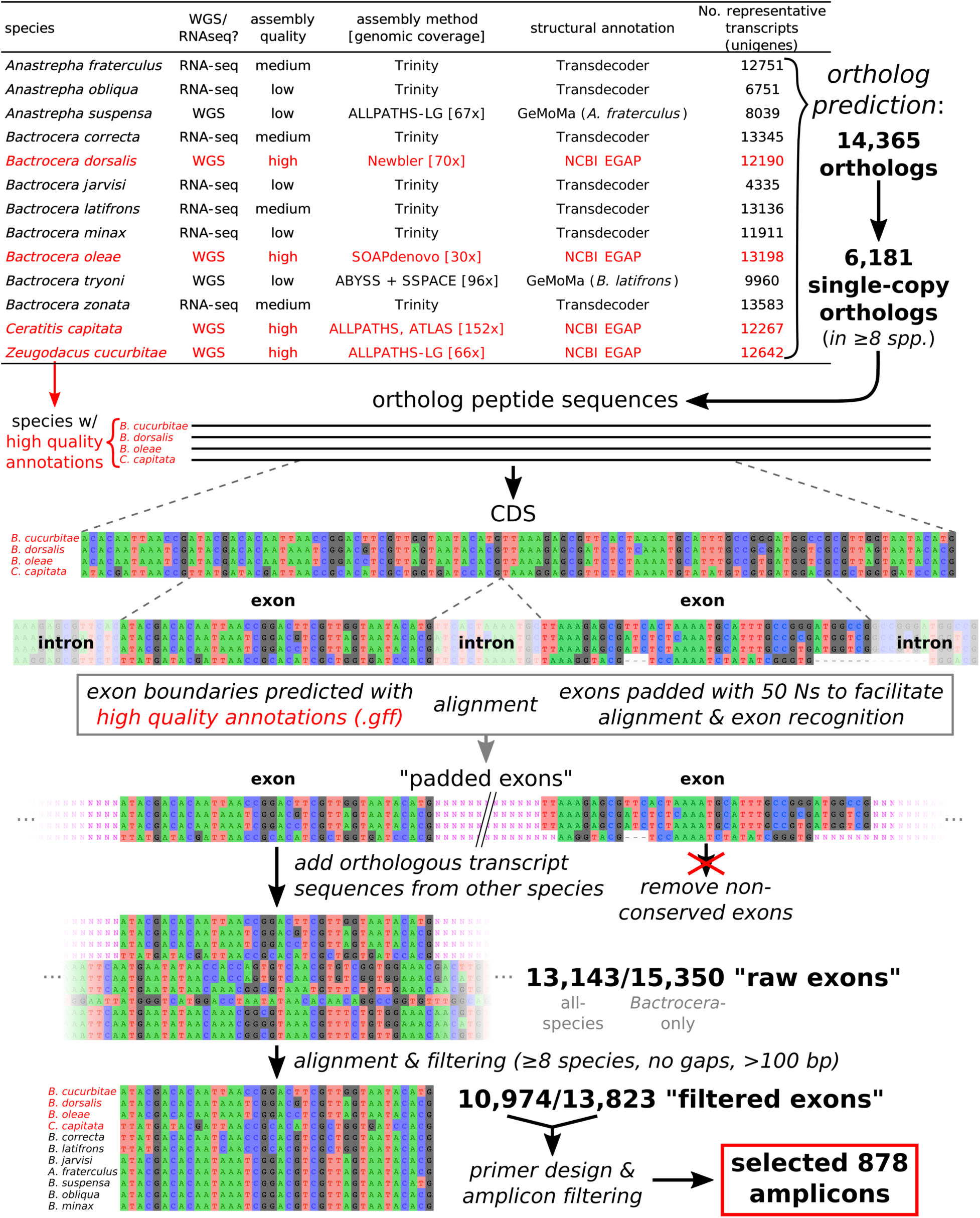
Visual depiction of bioinformatic locus selection pipeline. Processes/filtering steps are indicated with italics. Sequence visualizations generated with AliView v1.18 (Larsson 2014).

**Figure 2.**
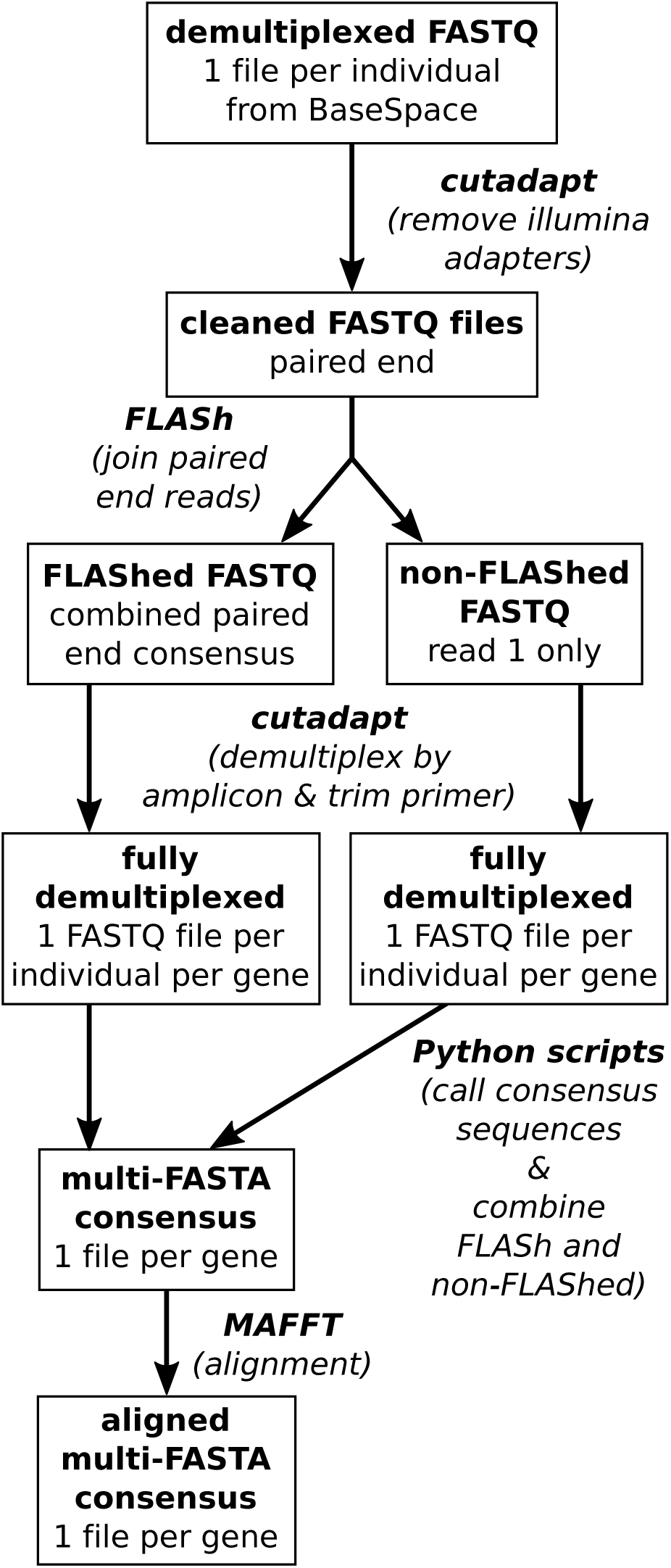
Flowchart of data processing steps. Processing/filtering steps are indicated with italics, and boxes indicate intermediate files.

### Locus Selection Pipeline

#### 1. Ortholog Prediction

Our locus selection pipeline begins with ortholog prediction from existing genomic and transcriptomic resources in the focal group. For this study, we used data from 13 species including *Anastrepha* (three species), *Bactrocera* (nine species), and *Ceratitis* (one species): *Anastrepha fraterculus* (Wiedemann 1830), *A. obliqua* (Macquart 1835), *A. suspensa* (Loew 1862), *Bactrocera correcta* (Bezzi 1916), *Zeugodacus cucurbitae* (Coquillet 1899), *B. dorsalis* (Hendel 1912), *B. jarvisi* (Tryon 1927), *B. latifrons* (Hendel 1912), *B. minax* (Enderlein 1920), B. *oleae* (Rossi 1790), *B. tryoni* (Froggatt 1897), *B. zonata* (Saunders 1841), and *Ceratitis capitata* (Wiedemann 1824). For three species (*A. obliqua*, *B. jarvisi*, and *B. minax*) we downloaded sequence read archives from the National Center for Biotechnology Information and conducted *de novo* transcriptome assembly using the trinity pipeline (Grabherr, *et al.* 2011), as described in Sim, *et al.* (2015). For two species (*A. suspensa* and *B. tryoni*) we used GeMoMa (Keilwagen, *et al.* 2016) with default parameters to predict gene models based on other phylogenetically proximate species’ transcriptomes (*A. fraterculus* and *B. latifrons*, respectively). The quality of these data sources ranged from that of chromosome-scale genome assemblies (*Z. cucurbitae*: Sim and Geib (2017)) and high quality transcriptomes (generated from all life stages, comprehensive filtering of erroneous transcripts/partial sequences, etc.: Calla, *et al.* (2014), Geib, *et al.* (2014), Sim, *et al.* (2015)) to genome assemblies from relatively low coverage resequencing experiments or tissue specific RNA-sequencing experiments (e.g. Rezende, *et al.* (2016), Fig. 1, details for all data sources in Table S1). Later steps in our locus selection pipeline depend on trustworthy genome-based annotations, specifically used to define exon/intron boundaries, so we identified four species (*B. dorsalis*, *B. oleae*, *C. capitata* and *Z. cucurbitae*) as having high quality genomic annotations based on the integration of homology-based and evidence-based predictions of gene function (e.g. Papanicolaou, *et al.* 2016, Sim and Geib 2017). We use “high quality annotations” to describe these four species in later steps.

For each gene locus for each species, we identified a single transcript with the longest open reading frame to be used as a representative protein sequence. We used OrthoMCL (Li, *et al.* 2003, Chen, *et al.* 2006, Chen, *et al.* 2007) with default parameters to predict orthologous groups using these longest representative proteins of each predicted gene across all species. We used `select_clusters_v2.pl` from Hahn, *et al.* (2014) to filter ortholog groups to those with a minimum of eight species that were single-copy (maximum median (`--max_median`) and mean number (`--max-mean`) of sequences per taxon = 1). In addition, orthologs had to be present in all of the four species with high quality annotations.

#### 2. Identifying Conserved Exons

From the filtered, single-copy orthologs and the corresponding genomes/transcriptomes (with fasta and generic feature format (gff) files), we developed a bioinformatic pipeline for target selection to identify conserved single-copy exons. This pipeline functions through a series of Python scripts and primarily uses Python v3.6 (Python Software Foundation 2017), BioPython (Cock, *et al.* 2009), SciPy (Jones, *et al.* 2001), NumPy (van der Walt, *et al.* 2011), gffutils (Dale 2013), Pandas (McKinney 2010), GNU parallel (Tange 2011), MAFFT (Katoh and Standley 2013), and TAPIR (Pond, *et al.* 2005, Faircloth, *et al.* 2012a). Using part01 of the HiMAP pipeline, nucleotide coding sequences (CDSs) corresponding to the single-copy orthologs were generated for each species based on their gff file. Each CDS consists of multiple exons, and at exon boundaries we removed the introns, which can vary dramatically in length between species, and replaced them with strings of 50 Ns. We refer to these concatenated exons and strings of 50 Ns as “padded exons”. The padded exons for these four species were then aligned using the default L-INS-i algorithm in MAFFT (Katoh and Standley 2013). The presence of these strings of 50 Ns helped to keep conserved exons together in the alignment (regardless of potentially widely variable intron sequence), and the script uses the stretches of 50 Ns to compare the start and stop coordinates of individual exons across the alignment. When all four species’ start and stop coordinates matched for an exon, that exon alignment (plus the 50 Ns on both the 5’ and 3’ ends) and the full unaligned, nucleotide ortholog sequence of the other species from the ortholog prediction process was written to a new file. During this step, multiple exon-specific files are potentially created from each ortholog file (e.g. multiple files would be created for gene-1_exon-A, gene-1_exon-B, gene-2_exon-A, gene-2_exon-B, etc.), and these are referred to as “raw exons”.

Raw exons were then aligned using the L-INS-i algorithm in MAFFT. Again, the padding with 50 Ns facilitated this alignment, as the alignment algorithm prefers to keep these blocks of Ns together. Preliminary alignments without this padding tended to erroneously break up the already aligned sequences of the four species with high quality annotations, due to sequence variation in the (potentially) longer sequences of the other species (which were full length ortholog sequences). Following this alignment, we filtered exons and discarded those that were < 100 bp long or contained gaps anywhere in their alignment. The remaining exons are presumably single copy and conserved in terms of length and exon/intron boundaries across all species, and served as the input for primer design steps. We refer to these as “filtered exons”.

#### 3. Putative Primer Design

We used Paragon Genomics’ CleanPlex Custom Panel Design Service (Paragon Genomics, San Francisco, CA, included in the CleanPlex Targeted Library Kit) to design primers for the filtered exons. This process accounts for standard primer selection parameters (primer size, melting temperature, etc.) as well as amplicon compatibility (interactions between primers for different amplicons), and we adapted the process in two ways. First, the filtered exons were highly variable (e.g. an average of 29.9% of bases were variable across the “all-species” filtered exons (see below), or qualitatively, every second to fourth base in most alignments), so we allowed for a single degenerate base to be included in each primer. Second, we wanted to target exonic regions that could be fully sequenced with a single primer pair utilizing paired-end 2 x 300 bp sequencing (rather than tiling multiple amplicons across an exon), so we set a maximum amplicon length, including the locus-specific primers, of 450 bp (compared to the previous maximum length of 300-350 bp). Minimum amplicon length was 125 bp.

To serve as an additional filtering metric, we used part02 of the HiMAP pipeline to calculate phylogenetic informativeness (PI) for each amplicon (here we refer to the amplified sequence, the sequence in between primers, as “amplicons”) using TAPIR v1.1 (Pond, *et al.* 2005, Faircloth, *et al.* 2012a), which implements the algorithm of Townsend (2007). TAPIR uses a reference tree to calculate PI, and for this tree we used a Maximum Likelihood (ML) consensus tree generated from a peptide alignment of the orthologs predicted by OrthoMCL. Again, we used `select_clusters_v2.pl` to filter orthologs to those that were single-copy and present in all species, which resulted in 490 orthologs. Each of the orthologs was aligned using MAFFT (L-INS-i), and the resulting fasta files were concatenated using `catfasta2phyml.pl` (Nylander 2016), generating an alignment of 302,549 peptides. A ML tree search was conducted in IQ-TREE v1.4.2 (Nguyen, *et al.* 2015) with 1,000 ultra-fast bootstrapping replicates (Minh, *et al.* 2013) and 1,000 replicates of the Shimodaira/Hasegawa approximate likelihood-ratio test (SH-aLRT: Guindon, *et al.* (2010)).

TAPIR requires a complete dataset (i.e. no missing individuals for any genes), however our exon filtering regime only required eight species present per exon. To accommodate this missing data in our single amplicon alignments, we used the topology generated in the aforementioned ML consensus tree to fill in any missing individual’s sequence with that of its closest relative. Although not ideal, this approach is realistic given the variable missingness of many phylogenetic datasets, and will only underestimate the PI of any given gene (having an identical sequence to close relatives provides no additional phylogenetic information for that clade), which is preferred to overestimating PI. With these complete amplicon alignments, we used TAPIR to calculate PI at six relatively evenly spaced points along the tree, based on branch length, and averaged those values to provide a single estimate of PI per amplicon.

#### 4. Final Amplicon Selection

The inclusion of three genera in this locus selection pipeline biases loci to those that are conserved at this relatively deep, genus-level phylogenetic scale. Given that our main focus and sampling effort was the genus *Bactrocera*, we wanted to maximize the phylogenetic resolution within the genus. For this reason, we ran our locus selection pipeline twice: once as described, starting with 13 species in three genera (“all-species”), and again with only *Bactrocera* species (“*Bactrocera*-only”). We used the same ortholog prediction results for both runs, so that ortholog IDs were conserved and we could avoid including duplicate exons in the final amplicon set. The *Bactrocera-*only procedure was identical to that with all species, except single copy orthologs were required to have five species to pass filter (instead of eight), three high quality annotations were used in the padded exon filtering, and filtered exons were required to have five species. We also used a minimum amplicon length of 175 bp, rather than 125 bp as in the all-species analysis.

With the two sets of amplicons (and putative primers), we selected our final amplicon set based on the following, in relative order of importance: roughly two-thirds of the amplicons being from the *Bactrocera*-only run and having no degenerate bases, product length (to maximize sequencing efficiency), PI, the presence of degenerate bases in the all-species primers, and the number of species from which orthologs were predicted. This was a relatively subjective process, and the first two criteria played the largest part in amplicon selection. PI cannot be compared between the all-species and *Bactrocera*-only runs, as a larger reference tree will automatically increase PI. We also only selected one exon per ortholog, as orthologs would generally be inherited as a single unit and share similar evolutionary history (however, multiple exons per ortholog could be targeted if longer loci were desired). Only when multiple exons (or the same exon from the two independent locus selection runs) had similar product length and relatively similar PI, did we consider the other criteria for amplicon selection. In other words, we only compared amplicon length, PI, the presence of degenerate bases, and the number of species (for which orthologs were predicted) when a single ortholog contained multiple potential amplicons; in this case we would compare these other attributes and select, for example, the longer amplicon or the one with the higher PI, subjectively. By avoiding tiling in locus design, and using a single amplicon per exon per ortholog, we aimed to maximize the unique loci sampled across the genome from a single amplicon pool, rather than maximizing locus length and sampling fewer unique loci. Tiling would also require subpooling of multiplex reactions, to avoid unintended amplicon products between primers in close proximity to each other. Primers were synthesized by a commercial vendor (Integrated DNA Technologies, IDT) and provided pooled in a single tube at a concentration of 250 nM.

### Specimen Collection and DNA Extraction

Specimens were collected as part of a larger effort to sample the diversity of fruit flies in the subfamily Dacinae. Adults were collected using male lure (cue-lure or methyl eugenol) or protein lure (Torula yeast) baited traps, as described in Leblanc, *et al.* (2013), and stored in ~95% ethanol in the field before being frozen at −80°C in the lab. Larvae were collected in their host fruits and reared to the adult stage. A total of 384 specimens were selected, and detailed collection information is provided in Table S2; most sampled flies belonged to *Bactrocera* and *Zeugodacus*, but we also included species of *Anastrepha*, *Ceratitis*, *Dacus*, *Neoceratitis*, and *Rhagoletis*. Whole flies were homogenized with 3.175 mm metal lysing beads, at a speed of 4.0 m/s for 20 seconds, in a FastPrep 24 homogenizer (MP Biomedical, Santa Ana, CA). The resulting homogenate was incubated at 55°C for three to twelve hours with proteinase K and tissue lysis buffer following manufacturer’s recommendations (Macherey-Nagel, Düren, Germany). We extracted DNA using a KingFisher Flex-96 automated extraction instrument (Thermo Scientific, Waltham, MA) and NucleoMag Tissue extraction kits (Macherey-Nagel, Düren, Germany), with an RNase A treatment, following manufacturer’s recommendations. DNA was eluted into 100 uL of Mag-Bind elution buffer, and quantified on a Fragment Analyzer Automated Capillary Electrophoresis System using a high sensitivity genomic DNA analysis kit (Advanced Analytical Technology, Ankeny, IA). We normalized DNA to 10 ng/µL using a Gilson PIPETMAX 268 (Gilson, Middleton, WI), unless the initial concentration was <10 ng/µL. The latter were left at their initial concentration. DNA quality was variable (Table S2) and some samples appeared to be of very poor quality or low concentration; we included some low-quality samples to test how the quality of input DNA impacts the resulting library.

### Library Preparation and Sequencing

We generated amplicons for all specimens using highly multiplexed PCR with a CleanPlex Targeted Library Kit (Paragon Genomics, San Francisco, CA). This library preparation consists of only three main steps and can be completed in <½ day of bench time: 1) amplify DNA targets using multiplex PCR, 2) digest non-specific products, and 3) add and amplify indexes and Illumina compatible adapters using a second PCR. Each of these steps is followed by purification using magnetic beads; for all purification steps we used in-house paramagnetic beads (as described in Rohland and Reich 2012) and an Alpaqua 96S Magnet plate (Alpaqua Engineering, Beverly, MA), and used a ratio of 1:3 sample:beads. We followed manufacturer’s recommendations for all library preparation steps. Briefly, for the first, locus-specific multiplex PCR, we used the 5X primer pool protocol, including 5 µL nuclease-free water, 2 µL 5X mPCR mix (Paragon Genomics), 2 µL 5X primer pool, and 30 ng (1 µL) of input DNA for samples that had an initial concentration >10 ng/µL or 50-60 ng (2 µL) for samples that had an initial concentration <10 ng/µL. Locus-specific primers used in the multiplex PCR had common sequences appended to their 5’ ends that matched the common sequence in the Illumina Nextera XT Index Kit v2 (i5 and i7, see below). Thermal cycling conditions consisted of a pre-heat step at 95°C for 10 minutes, 10 cycles of 98°C for 15 seconds followed by 60°C for 5 minutes, and a final hold at 10°C. To digest non-specific products, we added 6 µL of nuclease-free water and 2 µL each of the CP Reagent Buffer and Digestion Reagent to the cleaned PCR product, incubated at 37°C for 10 minutes, and used 2 µL stop buffer to terminate the reaction. In the second PCR, we added the Illumina Nextera XT Indexes (v2) by adding 18 µL nuclease-free water, 8 µL 5X 2^nd^ PCR Mix (Paragon Genomics), and 2 µL each of the i5 and i7 adapters (10 µM) to the cleaned, digested product. Using twenty-four i7 adapters and sixteen i5 adapters allowed individual indexing with 384 unique combinations. Thermal cycling conditions for this PCR were identical to the first PCR, except the 60°C portion of the cycle was held for 75 seconds, and eight cycles were used. The number of cycles was determined using manufacturer recommendations given 878 targeted amplicons and the generally low quantity of DNA.

We assessed library quality for each individual using a Fragment Analyzer, and a dsDNA 910 Kit (Advanced Analytical Technologies, Ankeny, IA), which is a qualitative kit that provides an accurate size distribution and a relative measure of sample concentration. Unfortunately, the Fragment Analyzer experienced mechanical difficulties during these runs and produced inconsistent results (see below), which were confirmed by quantifying several samples on a 2100 Bioanalyzer with the high sensitivity DNA Kit (Agilent, Santa Clara, CA). Given the low library volume at this stage (~6 µL), we used the Fragment Analyzer results to qualitatively bin samples into four categories: 1) “Good libraries” (good size distribution and relatively high DNA concentration), 2) “Moderate libraries” (good size distribution but lower DNA concentration), 3) “Poor libraries” (very low concentration, but size distribution still visible), and 4) “Blank libraries” (virtually no library present). Example traces from these subpools are provided in Figure S1. The fourth category of libraries was the most inconsistent between the Fragment Analyzer and Bioanalyzer, and in some cases a virtually nonexistent library as measured on the Fragment Analyzer was resolved on the Bioanalyzer. We pooled individuals from each of these library quality categories, and quantified these subpools using the Bioanalyzer as above. We then normalized subpools at equal molar ratios (considering the number of individuals per subpool) and generated a final library that was purified using paramagnetic beads (brought to a volume of 25 µL, using a ratio of 1:1 library:beads), and quantified using a Qubit 2.0 fluorometer with the dsDNA HS Assay Kit (Thermo Scientific, Waltham, MA). Paired-end 300 bp sequencing of the entire library was conducted on an Illumina MiSeq with sequencing reagent kit v3. Following preliminary analysis of this data, a second final normalized library was constructed from the moderate, poor, and blank libraries, and sequenced in the same fashion on a second run of the MiSeq sequencer; this second run was an effort to overcome the above errors in quantifying and normalizing the libraries.

### Data Processing

All data processing steps (summarized in Fig. 2) and phylogenetic analyses used default parameters and settings unless otherwise noted. Demultiplexing and FASTQ file generation was conducted using BaseSpace’s FASTQ Generation Analysis v1.0.0 (Illumina, San Diego, CA). From raw FASTQ files, we concatenated paired-end read files from both sequencing runs, removed Illumina adapters with cutadapt (Martin 2011), and used FLASh (Magoc and Salzberg 2011) to merge paired-end reads. We then used cutadapt to demultiplex each individual FASTQ file by amplicon by specifying each primer pair with the –`a` option, and requiring an overlap of 10 bp (`-O 10`). We then used part03 of the HiMAP pipeline to call a consensus sequence for each amplicon per individual. This script finds the most prevalent read length for each individual per amplicon and calls a degenerate consensus sequence based on all reads of that length, using the rules of Cavener (1987) via Bio.motifs in BioPython (Cock, *et al.* 2009). A minimum of five reads (per consensus) are required per individual per amplicon (this minimum read filter can be specified), and any consensus read that is <65 bp is removed (based on an observed natural break in the data). This script then calculates the mean length of the consensus sequences across all individuals per amplicon, and if an individual’s consensus sequence length deviates >20 bp from the mean (also based on an observed natural break in the data), it is removed. The output of this script is a single multi-FASTA per amplicon with consensus sequences for each individual using IUPAC ambiguities to represent heterozygous bases. For reads that could not be FLASh-merged (as identified by FLASh), we used the forward reads only (read 1), and demultiplexed and called consensus sequences as above. We then used an in-house Python script to compare the FLASh-merged and non-FLASh-merged consensus sequences, which added non-FLASh-merged sequences to the multi-FASTA files if those amplicon/individual combinations were not present in the FLASh-merged data. Finally, we incorporated the sequences from the data sources used for the locus selection pipeline (13 species), bringing the total number of taxa to 397.

Because loci were sequenced end-to-end, demultiplexing by primer sequences leads to individual consensus sequences that are already more-or-less aligned. However, some variation in sequence length within each amplicon was observed, generally due to biologically-real insertion/deletion events (3-9 bp differences). Therefore, we aligned all amplicons using the L-INS-I algorithm in MAFFT, before using `alignment_assessment_v1.py` (Portik, *et al.* 2016) and Bash scripts to assess data coverage across individuals and amplicons. We identified amplicon alignments with high proportions of gaps (>1%, 45 amplicons) and variable sites (>40%, 70 amplicons), as we observed that these characteristics were associated with off-target (non-matching) sequences and residual primer sequence. We checked these alignments manually using AliView v1.18 (Larsson 2014), removed offending individuals (from 11 amplicons total), and re-aligned before recalculating data coverage as above. Finally, we addressed missingness in the dataset by removing individuals that had <100 amplicons (< ~11% coverage), and amplicons that had <200 individuals (< ~52% coverage).

### Phylogenetic Analyses

We conducted two main types of phylogenetic analysis, general tree searches of concatenated datasets and species tree estimations, and used *Rhagoletis completa* Cresson 1929 as an outgroup for all analyses based on Segura, *et al.* (2007), Krosch, *et al.* (2012), and preliminary analyses. First, we conducted maximum likelihood (ML) and Bayesian inference (BI) based tree searches of the concatenated nucleotide alignment using IQ-TREE v1.4.2 (Nguyen, *et al.* 2015) and ExaBAYES v1.5 (Aberer, *et al.* 2014), respectively. IQ-TREE was run with 1,000 ultra-fast bootstrapping replicates (Minh, *et al.* 2013) and 1,000 replicates of the Shimodaira/Hasegawa approximate likelihood-ratio test (SH-aLRT: Guindon, *et al.* (2010)) to assess node support. The model of evolution was selected using IQ-TREE’s model selection procedure. For BI using ExaBAYES, four independent runs were conducted, each having four coupled chains and attempting four swaps per generation on average. Each independent run progressed for one million generations and was sampled every 1,000 generations. We assessed convergence and sampling of parameter values’ posterior distributions with Tracer v1.6 (Rambaut, *et al.* 2014), by ensuring that effective sample sizes were >200. We manually removed the first 25% of trees from each run as burn-in, combined post burn-in trees for all runs, and built a consensus tree using TreeAnnotator (Drummond, *et al.* 2012). Trees were visualized using FigTree v1.4.2 (Rambaut and Drummond 2010), GraPhlAn v0.9.7 (Asnicar, *et al.* 2015), and the APE library (Paradis, *et al.* 2004) in R v3.3.1 (R Core Team 2016).

We then tested two other main phylogenetic considerations using additional ML tree searches in IQ-TREE. First, we conducted partitioned analysis using a scheme estimated by PartitionFinder2 v2.1.1 (Lanfear, *et al.* 2017). We used the `rcluster` algorithm (with `rcluster-max` = 1,000 and `rcluster-percent` = 10, as suggested in the documentation) on all models for all nucleotide partitions (739 in total) with the AICc selection criteria in PartitionFinder2, and allowed partition-specific rates in IQ-TREE. Second, to accommodate for nucleotide saturation, we conducted analyses on peptide sequence alignments generated from the raw nucleotide alignments. To generate peptide sequences, we created a BLAST database from the concatenated input of the ortholog prediction for locus selection (the longest representative transcript of each predicted protein), and used BLASTX in BLAST+ (Camacho, *et al.* 2009) to predict peptide sequences from the nucleotide alignments. We set `–max_target_seqs` to one to output a single returned hit per individual per gene, and reformatted these outputs to fasta format before aligning with MAFFT, as before. We then manually checked alignments with gaps and those where the number of individuals did not match the number of individuals in the nucleotide alignment, and removed alignments where the BLASTX search was returning non-orthologous hits. Finally, as with the nucleotide datasets, we removed amplicons with <200 individuals and individuals with <100 amplicons. We conducted tree searches on the peptide-based alignment using IQ-TREE’s model selection procedure, as well as using a partitioning scheme estimated by PartitionFinder2, which was run in the same fashion as for the nucleotide alignment.

We estimated species trees from the nucleotide alignment using three methods: polymorphism-aware phylogenetic models (PoMo) (Schrempf, *et al.* 2016) in a ML framework, and coalescent-based frameworks with quartet inference in SVDquartets (Chifman and Kubatko 2014) and ASTRAL-II (Mirarab and Warnow 2015). These three methods are well-suited for this type of dataset because they allow for missing data among specimens and species, are amenable to large datasets, and can take into account multiple individuals per species (a parameter we enforced for all three analyses). Polymorphism-aware phylogenetic models incorporate population site frequency data to directly account for incomplete lineage sorting within species, and we implemented PoMo in IQ-TREE’s PoMo version v1.4.3 (Nguyen, *et al.* 2015). We used the `FastaToCounts.py` script in the cflib library (Schrempf, *et al.* 2016) to generate the allele counts input file, the general time-reversible (GTR) model with default PoMo additions, and 1,000 ultra-fast bootstrap replicates to assess node support. Quartet inference uses algebraic statistics to infer species relationships by considering the relationship among four species at a time (quartets), and summarizing quartets across the dataset (Chifman and Kubatko 2014). SVDquartets generates quartets from single-sites (single mutations) across a dataset, whereas ASTRAL-II takes individual gene trees as input and has been shown to outperform SVDquartets when incomplete lineage sorting is high (Chou, *et al.* 2015). We implemented SVDquartets in PAUP v4.a152 (Swofford 2017) and evaluated 10 million random quartets with *R. completa* set as the outgroup (preliminary analysis that evaluated all possible quartets produced virtually identical topologies). We generated gene trees for ASTRAL-II using the best trees generated by RAxML v8.2.4 (Stamatakis 2014) for each amplicon (using the GTRGAMMA model), and implemented the multiple individual version of ASTRAL-II v4.10.12 (Mirarab and Warnow 2015), using its default branch support measurement (Sayyari and Mirarab 2016). We quantitatively compared trees from both the main phylogenetic analyses and species tree analyses using normalized matching cluster distances (Bogdanowicz and Giaro 2013) in TreeCmp v1.1-b308 (Bogdanowicz, *et al.* 2012).

## RESULTS

### Locus Selection Pipeline

The 13 initial data sources for our locus selection pipeline consisted of six genomes and seven transcriptomes and had an average of 11,085 representative genes/unigenes per data source (Fig. 1). OrthoMCL predicted 14,365 orthologs, 6,181 of which were single-copy and present in at least eight species (including the four with “high quality annotations”). From these 6,181 orthologs, the two runs of our locus selection pipeline (all-species and *Bactrocera*-only, respectively) generated 13,143 and 15,350 raw exons and 10,974 and 13,823 filtered exons (see Methods for distinction). Given the anticipated number of reads per specimen generated from a single lane of sequencing on the MiSeq and read-depth, we aimed for between 800 and 1,000 amplicons (conservatively planning 20 million reads per run, 1000 amplicons, and 384 specimens equates to ~50x coverage). Thus, from the filtered exons for which primers could be made, we selected 372 and 730 primer sets (all-species and *Bactrocera*-only, respectively) to filter by amplicon characteristic (length, PI, presence of degenerate bases, etc.), and chose 878 amplicons for the final amplicon set. Primers were an average of 21.5 bp in length (without adapters for Illumina sequencing), and 205 contained degenerate bases.

### Sequencing and Data Processing

Two MiSeq runs produced 37.8 million reads (20.3 and 17.5, respectively) that were successfully demultiplexed by specimen, with an average of 98.6 thousand reads per individual. A total of 37.1 million reads were successfully demultiplexed by amplicon, with 35.4 million of those being FLASh-joined reads (95.4%), and the remaining 1.7 million being “read 1 only” reads (reads not successfully FLASh-joined, 4.6%). A total of 34.9 million reads were used to call consensus sequences across all individuals (average per individual: 99.4 thousand reads), which resolved a total of 227,499 individual consensus sequences out of a possible 337,152 (384 individuals x 878 loci). The average read depth per consensus sequence (N = 227,499) was 153.8 (±1.09). Only 394,085 reads (1.04% of all reads) matched to an individual and an amplicon, but were not used to call a consensus (i.e. off-target sequences).

An average of 592 amplicons (67.4%) were resolved per individual (Table 1). Thirty-two individuals (8.3%) failed, with no recovered amplicons, and an additional 17 individuals (4.4%) had poor performance with <100 recovered amplicons. An average of 259 individuals (67.4%) were resolved per amplicon, and <200 individuals were resolved in 139 amplicons (15.8%) (Table S3). Individuals in higher quality library subpools had more amplicons resolved (Table 1). However, we also observed individuals from the “blank” subpool (the lowest quality) that had high proportions of amplicons resolved (Fig. 3); this indicates that our original library QC was not accurate, but that this suboptimal QC did not bear directly on our end product (see Fig. S1). There was a modest trend for shorter amplicons (Fig. 4a), but all amplicons had comparable phylogenetic informativeness relative to amplicon length (Fig. 4b). Additionally, we observed no amplification bias based on whether the amplicon was selected from the all-species or *Bactrocera*-only locus selection pipeline, or whether degenerate bases were present in the primers (which were only allowed in primer design for the all-species amplicons) (Fig. 4a). Amplicon recovery was not related to initial DNA concentration (pre-library preparation: F_1,382_ = 0.04051, r^2^ = −0.002, *p* > 0.05), however these concentrations would not address inflation due to bacterial DNA contamination (a phenomenon we have observed with other specimens collected in this manner) or overall DNA quality (specifically fragment size). Finally, amplicon recovery did generally decrease with increased phylogenetic distance from *Bactrocera,* regardless of library subpool (Fig. 5).

**Table 1.** Sample sizes, and average (±95% confidence interval) raw reads, reads used to call consensus sequences, and recovered loci per subpool and overall (“all”). Statistics excluding failed or poor quality individuals indicated with asterisks: *excluding individuals with <100 genes, **excluding individuals with 0 genes.

| subpool | N | raw reads | consensus reads | recovered loci |
| --- | --- | --- | --- | --- |
| 1 | 38 | 87,441 ($\pm 8,298$) | 80,851 ($\pm 7,717$) | 740.1 ($\pm 34$) |
| 2 | 146 | 117,947 ($\pm 12,270$) | 109,202 ($\pm 11,509$) | 676.2 ($\pm 32.5$) |
| 2* | 144 | 119,535 ($\pm 12,240$) | 110,677 ($\pm 11,485$) | 684.8 ($\pm 30.7$) |
| 3 | 113 | 97,190 ($\pm 15,821$) | 92,887 ($\pm 14,794$) | 610.3 ($\pm 45.2$) |
| 2** | 109 | 100,754 ($\pm 16,011$) | 92,887 ($\pm 14,794$) | 632.7 ($\pm 41.2$) |
| 3* | 103 | 106,482 ($\pm 12,859$) | 98,177 ($\pm 15,036$) | 666.9 ($\pm 33.1$) |
| 4 | 87 | 73,171 ($\pm 36,129$) | 99,143 ($\pm 47,468$) | 363.7 ($\pm 70.8$) |
| 4** | 59 | 107,831 ($\pm 44,083$) | 99,143 ($\pm 47,468$) | 536.3 ($\pm 69.6$) |
| 4* | 50 | 127,110 ($\pm 58,736$) | 116,941 ($\pm 54,630$) | 627.9 ($\pm 49.1$) |
| all | 384 | 98675 ($\pm 10644$) | 99403 ($\pm 10374$) | 592.4 ($\pm 27.6$) |

**Figure 3.**
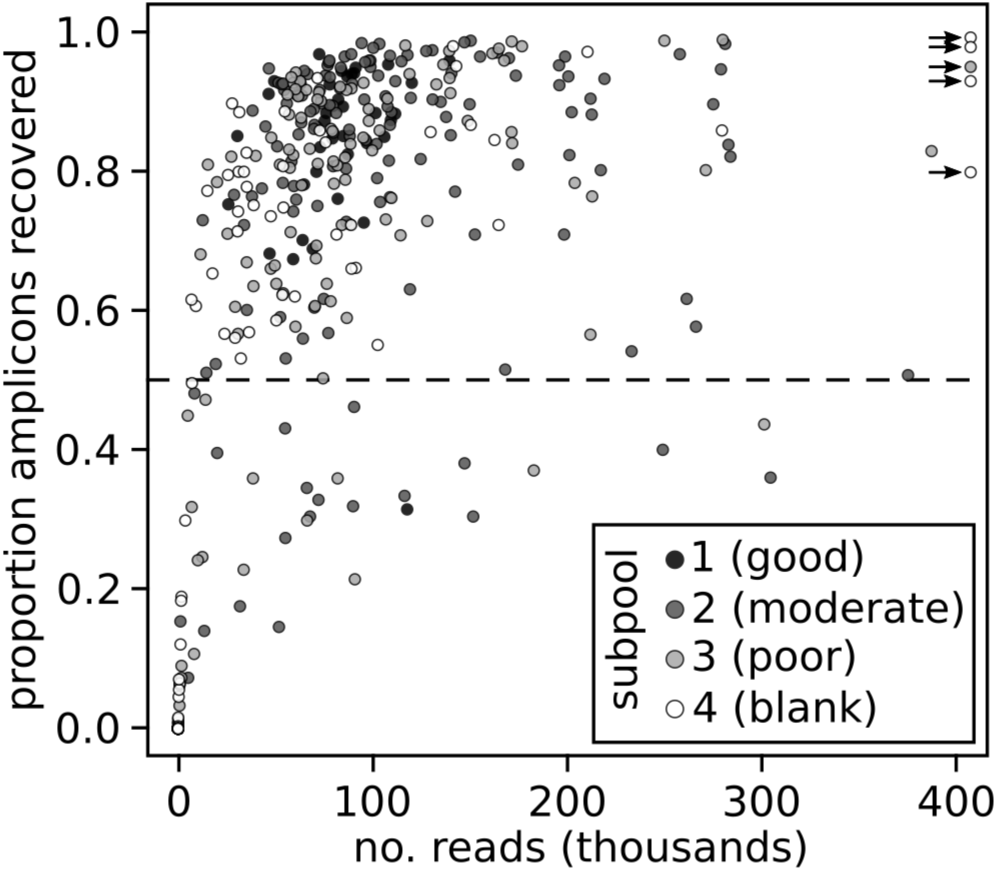
Proportion of amplicons recovered (out of 878) versus the number of reads per individual.

**Figure 4.**
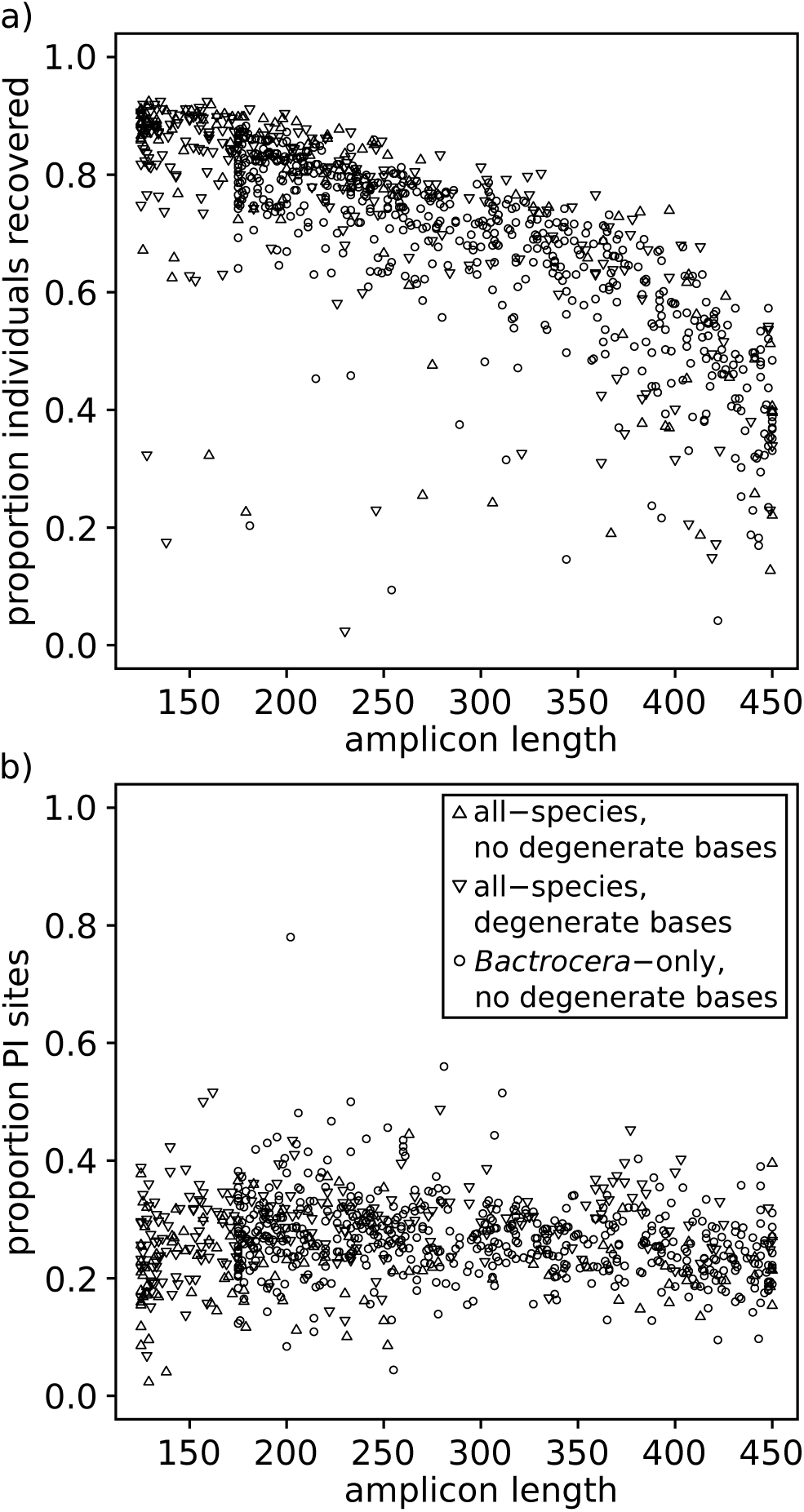
a) Proportion of individuals recovered (out of 384) versus amplicon length, and b) proportion of Parsimony Informative (Par. Inf.) sites per amplicon versus amplicon length. The shape of each amplicons is according to whether they were developed for all-species or *Bactrocera*-only, and if their primers contained degenerate bases.

**Figure 5.**
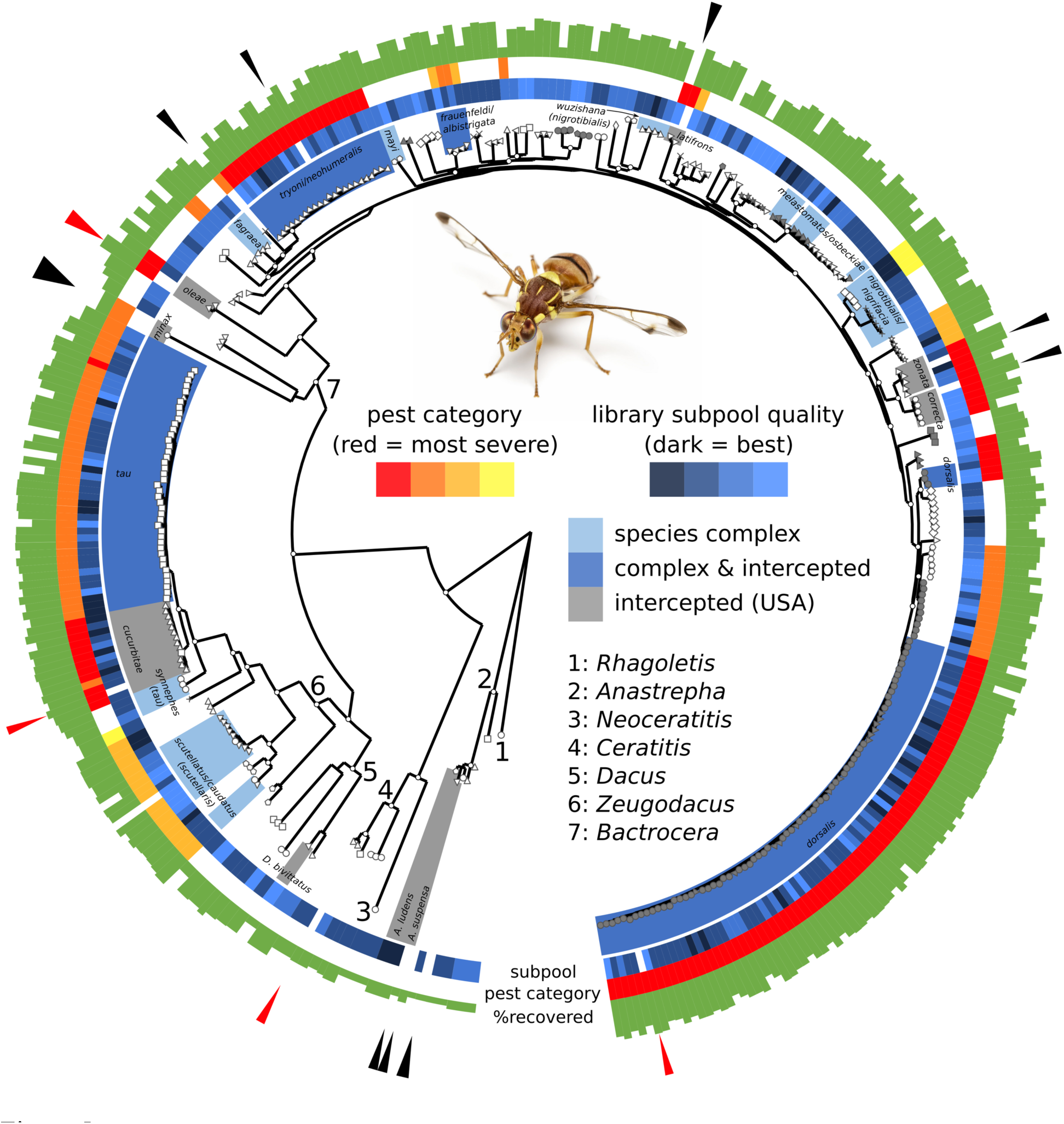
Consensus tree from Maximum Likelihood analysis of 739 conserved exons in peptide space (49,528 amino acids in concatenated alignment) for 348 individuals. Genus-level and higher classifications are denoted with numbers on nodes, and main nodes that are highly supported (SH-aLRT/ultra-fast bootstrap >0.8/0.95) are labeled with white circles (for intraspecific node support, see Fig. S5). Shapes at terminal branches indicate species identifications, and species that belong to historically difficult species complexes or are intercepted in the United States of America, or both, are indicated with boxes around the clades and species epithets/complex names (parentheses following species epithet indicates the complex in which that species belongs). The *B. dorsalis* complex is the only exception: here, species now synonomized to *B. dorsalis* are indicated with a box around the clade, other members of the *B. dorsalis* complex are indicated with grey shapes at terminal branches. Library subpool quality (“subpool”, see Methods), pest category based on Vargas, *et al.* (2015) (“pest category”), and the percent recovered loci (“%recovered”, out of 878) are displayed on rings around the tree, and arrows outside these rings indicate specimens whose data were generated from the data sources used in the locus selection pipeline (red arrows indicate “high-quality” annotations). Inset photograph of *Zeugodacus cucurbitae* by A.N. Suresh Kumar, used with permission.

### Phylogenetic Analyses

Our final filtering to address missingness in the nucleotide alignments removed 49 individuals and 138 amplicons, leading to a final dataset of 739 amplicons (151,511 bp concatenated alignment) for 348 individuals. Similar filtering for the peptide-based alignments resulted in 734 amplicons (49,528 peptides concatenated alignment) for 348 individuals. Phylogenetic analyses generally produced similar main topologies regardless of method. Figure 5 shows a representative topology generated with ML of the peptide alignment, and all trees (including models and partitioning statistics) are provided in Figures S2-S6. All methods agreed on the main relationships between genera included here: *Anastrepha*, *Ceratitis*, *Neoceratitis, Dacus*, *Zeugodacus*, and *Bactrocera*. These relationships included a sister relationship between *Dacus* and *Zeugodacus*, which has received mixed support in previous studies (Virgilio, *et al.* 2015). Most species were reciprocally monophyletic, except in the cases of known complexes consisting of morphologically similar species (Fig. 5). A small number of individuals were placed in unexpected positions on the tree (noted in Fig. S2). In some cases, this appeared to be the result of potential specimen misidentifications or a mix-up during specimen or library preparation, for example one *Z. cucurbitae* and one *Z. tau* individual being placed in the other’s respective clade. In other placements, it is less clear whether misidentifications or biological causes (cryptic species) are to blame, as in the case of *B. fuscitibia* being placed in disparate clades on most trees (e.g. Fig. S2). Node support was generally high across the tree regardless of method (SH-aLRT/ultra-fast bootstrap > 0.8/0.95 or posterior probability > 0.9), and qualitatively, terminal branch lengths were slightly longer in partitioned analysis and the peptide-based analysis.

The main discordance between methods was observed in relationships between groups of closely-related *Bactrocera* species. This discordance is most easily visualized when comparing the species tree estimations (Fig. 6), although similar discordance was observed when comparing consensus trees from ML and BI analyses as well as nucleotide and peptide alignments (Figs. S2-S6). Most sister species pairs and complexes were conserved across analyses (e.g. (*B. nigrifacia, B. nigrotibialis*) and (*B. unirufa*, (*B. wuzishana*, *B. amplexiseta*))), however, the mid-level relationships between these groups were more variable, and had lower node support in all analyses. Regardless, pair-wise normalized matching cluster distances between trees from both the main phylogenetic analyses (all specimens) and the species tree analyses indicated that overall these trees were quite similar to each other (Table S4). The potential species misidentifications mentioned above could impact species tree estimations, particularly when few specimens are sampled per species. However, the similar discordance between general tree searches (multiple specimens per species) and species tree methods suggest that the discordance observed may be more likely a result of data characteristics (genes having different evolutionary histories) rather than potential misidentifications.

**Figure 6.**
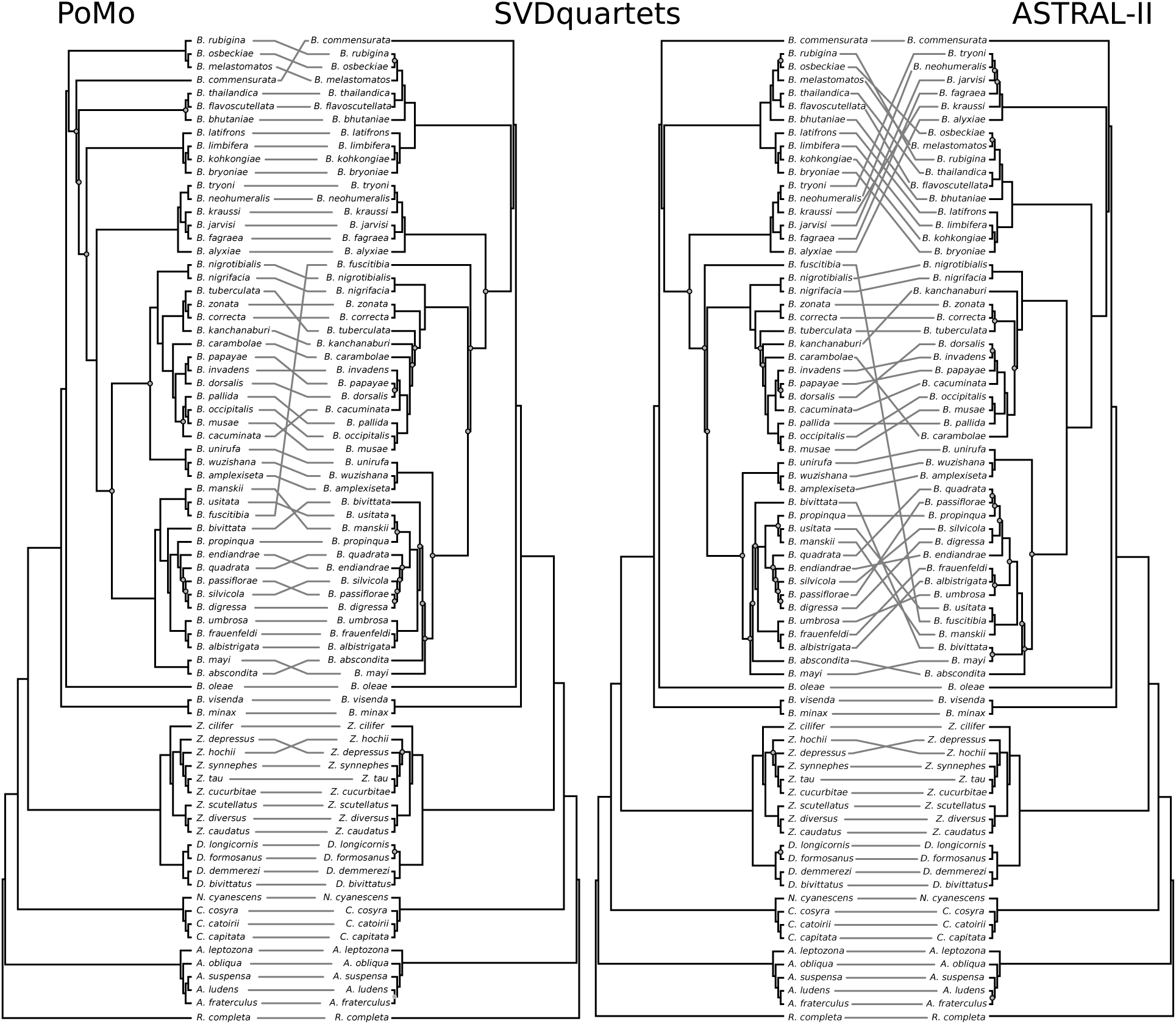
Comparison of species trees estimated with three methods: Polymorphism-aware phylogenetic model to compared to SVDquartets (left), and SVDquartets compared to ASTRAL-II (right). To facilitate comparison, the SVDquartets tree in each comparison is identical. Grey dots on nodes indicate bootstrap support <90% in each respective method.

## DISCUSSION

Here we demonstrate HiMAP, a new approach for building phylogenomic datasets using highly multiplexed amplicon sequencing. This methodology is relatively inexpensive and easily amenable to large numbers (hundreds) of taxa, requires minimal hands-on time at the bench, and data can be processed rapidly for consensus calls, avoiding read mapping or assembly. We discuss the advantages and disadvantages of HiMAP, our locus selection pipeline, and briefly, systematic conclusions from this dataset.

### Phylogenomic Data Collection

Choice of phylogenomic data collection method often boils down to a few main logistical consideration and trade-offs (reviewed in Lemmon and Lemmon 2013), and the most commonly-used methods have clear strengths and limitations. HiMAP, combined with current advances in amplicon sequencing technology, provides an alternative strategy for cost-effective phylogenomic datasets of moderate to long length loci. Cost is a main consideration for any genomic study, and one of HiMAP’s strengths is its relatively low cost, with a per-specimen cost of $40-50 USD/specimen (list prices for this dataset provided in Table S5). This price is predominately dependent on three relatively expensive items, the library preparation kit, primers, and sequencing, but is roughly on-par with estimates for the per-specimen cost of sequencing UCEs (~$65/specimen: Faircloth, BC, *personal communication*). Additional advantages of this approach are that amplicon library preparation is simple (it consists of two PCR reactions per sample, a digestion step, and intermediate clean up steps) and fast (2.5 hours for a handful of specimens, a half day for 48-96 specimens at a time), equating to low personnel costs. Finally, this approach generates very efficient use of sequencing reads (92% of the raw reads were used to call consensus sequences). Taken together, these characteristics lead to high overall cost-effectiveness.

The challenges of multiplex PCR generally have revolved around issues of substantial validation and optimization of multiplex reactions, inconsistent and off-target amplification, limited utility as phylogenetic distance increases, and the prevalence of primer dimers (Markoulatos, *et al.* 2002, Lemmon and Lemmon 2013). Ultimately, given the advances in current amplicon sequencing technology, we experienced very few of these challenges. We conducted no validation or optimization of multiplex reactions other than filtering out exons that had primer compatibility issues, and had very few cases of off-target amplification (most of which were removed automatically in data filtering (see below); only 11 amplicon alignments were manually edited to remove off-target sequences). By allowing a single degenerate base per primer we aimed to broaden the phylogenetic utility of this amplicon set, and we were able to recover hundreds of loci in genera separated by 65-100 million years (*Bactrocera* vs. *Rhagoletis* or *Ceratitis*: Krosch, *et al.* (2012), Tephritidae: Caravas and Friedrich (2013)). The amplification procedure and proprietary digestion step of the CleanPlex Targeted Library Kit were effective in limiting and removing primer/adapter dimers, respectively, as we observed no signs of dimers during final library preparation QC or in sequencing results (Fig. S1). Input DNA requirements are quite low (tens of nanograms), and we observed high amplification success in samples with low DNA quality. Additionally, although our library QC was less than ideal, we were able to achieve high amplicon recovery (>80%) from samples across a broad range of quality and source, including those with low DNA quantity (Table S2). A simpler library QC process, involving library quantification rather than measuring the library size distribution, with spot checks of size distribution as we did here, may be more effective than methods used in this study. Although further research will be required to validate the minimum required DNA quality and quantity, our results are optimistic that this approach may work reasonably well with historic or museum specimens. The CleanPlex library technology was designed to be implemented with cancer tissue samples, or other biopsy tissue that may be preserved as formalin-fixed paraffin-embedded (FFPE) specimens; these DNA sources often have similar quality issues as historical museum samples (low quality and quality). Additionally, tiling of smaller amplicons (100-200 bp) across a target region could facilitate use of this method with historical samples, and would provide a useful comparison with other phylogenomic approaches using museum specimens (Blaimer, *et al.* 2016b, McCormack, *et al.* 2016).

Finally, data processing from raw sequence data to consensus sequences, like the wet lab steps, is streamlined and fast. The primary reason for this is that locus recovery is ideally tailored to be end-to-end, so that data processing can be assembly-free (a maximum amplicon length of 450 bp was used here to accommodate 2x300 bp sequencing). Our primary data processing approach consists only of two steps of demultiplexing with cutadapt (Martin 2011), merging paired-end reads with FLASh (Magoc and Salzberg 2011), DNA alignment with MAFFT (Katoh and Standley 2013), and filtering using a custom Python script (Fig. 2). This approach is analytically straightforward, and could theoretically be accomplished on a modern laptop in a reasonable amount of time; with the use of modest high-performance computing resources, all data processing steps for a dataset of the size presented here can be conducted in several hours. Additionally, the standard output format (aligned multi-FASTA format) is easily used directly, or with simple reformatting, by many routine phylogenetic software packages. This consensus sequence approach streamlines the transition from data processing to phylogenetic analysis, although other more analytically intensive procedures (e.g. phasing, discussed below) might be favorable.

Overall, HiMAP provides a cost-effective strategy to generate moderate to long length phylogenomic loci for hundreds of individuals in a time-efficient manner. Compared to other phylogenomic data collection approaches, the overall alignment length of this dataset is similar to some of those generated with AHE or UCE approaches (e.g. Brandley, *et al.* 2015, Breinholt, *et al.* 2017), and the ability to efficiently sequence hundreds of specimens is advantageous. Including more species and more specimens per species is an important consideration, as adding additional taxa, even with high missing data, has been shown to increase phylogenetic accuracy (Wiens and Tiu 2012) and reduce nonphylogenetic signal caused by systematic error (Baurain, *et al.* 2007, Philippe and Roure 2011). Additionally, the ability to target specific loci for individual experiments in a cost-effective manner, rather than relying on a single locus- or probe-set generated for a broad taxonomic group, is an attractive characteristic. In this way, locus selection can be tailored for particular research questions and phylogenetic scales, and thus be more efficient in targeting highly informative loci for each study.

### Locus Selection Pipeline & Data Processing

One limitation of this overall method is that it does require a set of previously selected loci from which to develop primers, although many other commonly-used approaches share this limitation (Lemmon and Lemmon 2013, Jones and Good 2016). To this end, we developed a bioinformatic locus selection pipeline that ingests virtually any genomic or transcriptomic resource, and predicts conserved, orthologous exons that are also phylogenetically informative. One or more “high-quality” annotation(s) is required by this pipeline to predict exon boundaries across all data sources, but besides this high-quality annotation, data sources of any quality can be used. This provides a valuable use for relatively low quality data sources (preliminary genome sequencing or resequencing experiments, tissue-specific RNA-Seq experiments, etc.) that may be difficult to use for other genomic endeavors. With the steadily decreasing cost of high throughput sequencing, the availability of genomes and transcriptomes should continue to increase, thus providing valuable data for comparative applications such as ours here (see also Faircloth 2017).

In this study we targeted relatively short amplicons, compared to many probe-based approaches that often target longer genes through tiling of multiple probes across a gene (Faircloth, *et al.* 2013, Breinholt, *et al.* 2017). Our goal was to sample as many unlinked genomic regions as possible, rather than fewer, but longer, loci. We found no relationship between amplicon length and its relative phylogenetic informativeness (Fig. 4b), suggesting that the results of this approach are not biased by targeting shorter amplicons. While data processing is simplified by sequencing amplicons in an end-to-end fashion, it also places a limit on amplicon length that is absent in sequence capture approaches. However, if longer length amplicons were desired (e.g. to increase resolution of gene tree analyses), multiple exons per gene could be targeted as individual amplicons, and concatenated end-to-end data processing.

We envision several ways to increase the robustness of the data processing pipeline and analysis of HiMAP data. First, we are solely relying on the ortholog prediction process (and in turn the quality of the genomic/transcriptomic inputs to that process) and general data alignment and filtering to eliminate potential paralogs in our dataset. While we are confident in the overall quality of most of these inputs, particularly the “high quality” ones used extensively in the locus selection pipeline, additional validation steps could be used to ensure that the final datasets consist only of single-copy orthologs (e.g. Kristensen, *et al.* 2011). Second, this data processing approach does not account for PCR duplicates. Unlike many next-generation sequencing libraries, there is no random shearing step which facilitates the identification of PCR duplicates. Here, the only post-hoc method of avoiding PCR duplicates would be to increase minimum read depth for consensus calling; however, given the very high average read depth, setting this threshold would be an arbitrary decision (preliminary analyses with a higher read depth resulted in similar phylogenetic conclusions). The use of a “molecular tag” or “unique molecular identifier”, which is short randomer appended to the sequencing primers (Kinde, *et al.* 2011, Yourstone, *et al.* 2014, Kou, *et al.* 2016), would facilitate identification and removal of PCR duplicates. We recommend such an approach for future HiMAP studies, to quantitatively evaluate the effect of PCR duplicates, and investigate appropriate minimum read-depth for this type of data. Finally, by calling a consensus sequence for heterozygote base calls, rather than phasing the diploid sequence data into haplotypes, we are potentially losing valuable phylogenetic information (Browning and Browning 2011). Although beyond the scope of this study, implementing haplotype phasing into the HiMAP pipeline, or downstream analyses, would be of great interest.

### Systematic Conclusions

To date, molecular phylogenetic studies of Tephritidae have been limited to traditional molecular systematic approaches (<10 genes); the concatenated nucleotide alignment generated here is >50 fold larger than the most recent comparable dataset (Virgilio, *et al.* 2015). Although the current study also sequenced more specimens than previous studies, we focused on sequencing multiple specimens per species and thus sampled fewer species overall (Krosch, *et al.* 2012, Virgilio, *et al.* 2015). For this reason, we limit our systematic conclusions to those dealing with generic relationships and morphologically-difficult species complexes for which we had intensive sampling.

All phylogenetic analyses agreed on a single set of relationships between genera within the subtribe Dacina (*Bactrocera*, *Dacus*, and *Zeugodacus*), and notably supported a sister relationship between *Dacus* and *Zeugodacus*. Virgilio, *et al.* (2015) recently elevated *Zeugodacus,* formerly a subgenus of *Bactrocera* (*Zeugodacus*), to the generic rank with data from four mitochondrial genes and a single nuclear gene. Although the authors suspected a sister relationship between *Dacus* and *Zeugodacus*, which would support earlier conclusions of Krosch, *et al.* (2012) again based on a handful of mitochondrial and nuclear genes, the exact relationships between genera within Dacina were unclear. Our dataset strongly supports this relationship, as well as the generic status of *Zeugodacus*.

We focused a large portion of our sampling on morphologically-challenging species complexes within *Bactrocera* and *Zeugodacus* (Fig. 5). Some of these complexes have received substantial taxonomic and systematic attention, such as the *B. dorsalis* complex, where the synonymization of several species (Schutze, *et al.* 2015) has been argued (Drew and Romig 2016, Schutze, *et al.* 2017). Here, we asked whether our methods could distinguish between members of these complexes or not (i.e. reciprocal monophyly vs. para-/polyphyly). In some cases, we did find reciprocally monophyletic relationships between members of the complex (*Z. tau* & *Z. synnephes* and the *B. nigrotibialis* complex), but in most we were unable to fully separate members of the complex (the *Z. scutellaris*, *B. tryoni*, and *B. frauenfeldi* complexes) (Fig. 5). The *B. dorsalis* complex consists of a large number of morphologically and ecologically similar species, including several that have been synonymized with *B. dorsalis* (Drew and Romig 2013, Schutze, *et al.* 2015). We found support for the recent synonymization by Schutze, *et al.* (2015) (for clarity when referencing this synonymization, we use the pre-synonymization names in all supplementary trees), but otherwise find that most members of this complex are distinct and phylogenetically disparate from *B. dorsalis sec.* Leblanc, *et al.* (2015) and Schutze, *et al.* (2015). It is clear from this phylogenomic dataset that more detailed sampling and treatment of these complexes, on an individual basis, will be required to elucidate their evolutionary histories and potentially reevaluate their taxonomy. Additionally, more thorough sampling of species across the subtribe Dacina will be needed to evaluate general phylogenetic relationships within *Bactrocera*, *Dacus*, and *Zeugodacus*. The genome-wide markers developed here were selected based on their phylogenetic informativeness, and they should serve as a springboard for future genomics research in the Tephritidae.

### Conclusions and Prospectives

Here we present HiMAP, a novel approach for generating phylogenomic datasets using highly multiplexed amplicon sequencing. Both the wet lab and data processing components are rapid and straightforward, and the overall approach generates inexpensive datasets of hundreds to thousands of genes for several hundred individuals. Given its unique strengths compared to other phylogenomic data collection methods, we hope this study serves as a foundation to further develop this approach. We envision several main ways to increase overall efficiency and cost-effectiveness. First, multiplexing could potentially be increased substantially, as the CleanPlex Targeted Library technology has been used to multiplex up to 20,000 reactions in a single tube. However, as the number of targets increase, the cost of oligonucleotide synthesis also increases linearly, which may decrease the overall cost-effectiveness. Despite the linear cost increase, the oligonucleotides themselves are simple, unmodified oligos with no costly base modifications needed or additional purification, so could be generated through different methods. Alternative methods for oligonucleotide synthesis (e.g. array-based synthesis) may provide a more cost-effective way to increase multiplexing potential, however accounting for low yields using these approaches may prove to be a challenge. Additionally, multiplexed amplicon PCR technologies are rapidly improving and multiple providers are now creating such library kits; aspects of the HiMAP concept could be applied to various multiplex library preparation technologies (e.g. the Illumina TruSeq Custom Amplicon approach using extension ligation) or even alternative sequencing platforms (e.g. Thermo Fisher Ion AmpliSeq Panels using an Ion Torrent platform). Second, we could work to maximize the multiplexing of individuals in a run. We likely far exceeded required depth for many of our samples, with mean amplicon coverage of >100x, suggesting more individuals could have been indexed per library in this sequencing run. Optimizing the evenness of sample loading (through more accurate library QC, or working with standardized DNA input rather than variably low quality samples) would provide even greater potential for maximizing the multiplexing of a sequencing run, and increase overall efficiency.

Third, the MiSeq sequencing used here is on the low end of the sequencing output spectrum; targeting slightly shorter loci (200-300 bp) and sequencing on a HiSeq platform would greatly increase overall sequencing depth. Increasing sequencing output would additionally facilitate the pooling of more individuals and loci into a sequencing run. Extending the approach in such a way will require careful calculation of the balance between the number of targets, the number of individuals, anticipated sequencing depth, desired locus length, and cost. Considering the maximum output per platform, a single lane of sequencing on a HiSeq4000 produces 25x more data than a single run on a MiSeq, thus translating to the potential to sequence, for example, 5,000 amplicons for 1,024 individuals with >100x coverage (and accommodating for lower sequencing output). Sequencing on a HiSeq platform would also facilitate comparison with other methods (AHE, UCE, etc.), as these methods most often use the HiSeq platform. Finally, we focused on a relatively conservative phylogenetic scale here, and it will be important to test this method’s limits with regard to phylogenetic divergence. This will be relatively specific to each primer set, but general trends may emerge with in-depth exploration. Ultimately, we hope this study provides a starting point to further develop HiMAP, and continue to explore global biodiversity through the lens of genomics.

## ACKNOWLEDGEMENTS

We thank Ivy Wan and Shaobin Hou for assistance with sequencing that was conducted at the Advanced Studies in Genomics, Proteomics and Bioinformatics core facility at the University of Hawai’i at Mānoa, Boyd Mori for statistical assistance, Nicole Yoneishi and Jaymie Masuda for lab work, and Edward Braun and Brant Faircloth for their insightful comments on this manuscript. We thank Bishnu Bhandari, Kemo Badji, J. Caballero, Salley Cowen, Elaida Fiegalan, M. Aftab Hossain, Chia-Lung Huang, H.Y. Huang, David Haymer, Will Haines, Y.F. Hsu, Akito Kawahara, Sada Lal, Yuchi Lin, R. Messing, Aiko Ota, Sylvain Ouedrago, Rudolph Putoa, N. Pierce, J. Quintana, Eric Rodriguez, T. Stark, Ema Tora Vueti, Misael Valladares, L.H. Want, James Walker, Koon Hui Wang, Tianlin Xian, and APHIS technicians for collecting specimens. Funding for this project was provided by United States Department of Agriculture (USDA) Animal and Plant Health Inspection Service (APHIS) Farm Bill Section 10007 projects “Diagnostic Resources to Support Fruit Fly Exclusion and Eradication, 2012-2014” and “Genomic approaches to fruit fly exclusion and pathway analysis, 2015-2016” to USDA-APHIS, USDA-ARS and UH Manoa (projects 3.0251.02 and 3.01251.03 (FY 2014), 3.0256.01 and 3.0256.02 (FY 2015), and 3.0392.02 and 3.0392.03 (FY 2016)). Bioinformatic tools and source code available at https://github.com/popphylotools/HiMAP, and raw sequencing data files are available at NCBI BioProject PRJNA398162 and SRA SRR5976662-SRR5977045; a stable release of all bioinformatic tools will be available from the Dryad Digital Repository. Figures were created using R (R Core Team 2016), Inkscape v0.91 (The Inkscape Team 2017), and GraPhlAn v0.9.7 (Asnicar, *et al.* 2015). USDA is an equal opportunity employer. Mention of trade names or commercial products in this publication is solely for the purpose of providing specific information and does not imply recommendation or endorsement by the U.S. Department of Agriculture.

## REFERENCES

Aberer AJ, Kobert K, Stamatakis A. 2014. ExaBayes: massively parallel bayesian tree inference for the whole-genome era. Mol Biol Evol, 31:2553–2556.

Asnicar F, Weingart G, Tickle TL, Huttenhower C, Segata N. 2015. Compact graphical representation of phylogenetic data and metadata with GraPhlAn. PeerJ, 3:e1029.

Barrow LN, Ralicki HF, Emme SA, Lemmon EM. 2014. Species tree estimation of North American chorus frogs (Hylidae: Pseudacris) with parallel tagged amplicon sequencing. Mol Phylogenet Evol, 75:78–90.

Baurain D, Brinkmann H, Philippe H. 2007. Lack of resolution in the animal phylogeny: closely spaced cladogeneses or undetected systematic errors? Mol Biol Evol, 24:6–9.

Blaimer BB, Brady SG, Schultz TR, Lloyd MW, Fisher BL, Ward PS. 2015. Phylogenomic methods outperform traditional multi-locus approaches in resolving deep evolutionary history: a case study of formicine ants. BMC Evol Biol, 15:271.

Blaimer BB, LaPolla JS, Branstetter MG, Lloyd MW, Brady SG. 2016a. Phylogenomics, biogeography and diversification of obligate mealybug-tending ants in the genus Acropyga. Mol Phylogenet Evol, 102:20–29.

Blaimer BB, Lloyd MW, Guillory WX, Brady SG. 2016b. Sequence Capture and Phylogenetic Utility of Genomic Ultraconserved Elements Obtained from Pinned Insect Specimens. PLoS One, 11:e0161531.

Bogdanowicz D, Giaro K. 2013. On a matching distance between rooted phylogenetic trees. International Journal of Applied Mathematics and Computer Science, 23.

Bogdanowicz D, Giaro K, Wrobel B. 2012. TreeCmp: Comparison of Trees in Polynomial Time. Evolutionary Bioinformatics:475.

Brandley MC, Bragg JG, Singhal S, Chapple DG, Jennings CK, Lemmon AR, Lemmon EM, Thompson MB, Moritz C. 2015. Evaluating the performance of anchored hybrid enrichment at the tips of the tree of life: a phylogenetic analysis of Australian Eugongylus group scincid lizards. BMC Evol Biol, 15:62.

Branstetter MG, Danforth BN, Pitts JP, Faircloth BC, Ward PS, Buffington ML, Gates MW, Kula RR, Brady SG. 2017a. Phylogenomic Insights into the Evolution of Stinging Wasps and the Origins of Ants and Bees. Curr Biol, 27:1019–1025.

Branstetter MG, Jesovnik A, Sosa-Calvo J, Lloyd MW, Faircloth BC, Brady SG, Schultz TR. 2017b. Dry habitats were crucibles of domestication in the evolution of agriculture in ants. Proc Biol Sci, 284.

Breinholt JW, Earl C, Lemmon AR, Lemmon EM, Xiao L, Kawahara AY. 2017. Resolving relationships among the megadiverse butterflies and moths with a novel pipeline for Anchored Phylogenomics. Systematic Biology, doi:10.1093/sysbio/syx048.

Browning SR, Browning BL. 2011. Haplotype phasing: existing methods and new developments. Nat Rev Genet, 12:703–714.

Calla B, Hall B, Hou S, Geib SM. 2014. A genomic perspective to assessing quality of mass reared SIT flies used in Mediterranean fruit fly (Ceratitis capitata) eradication in California. BMC Genomics, 15:98.

Camacho C, Coulouris G, Avagyan V, Ma N, Papadopoulos J, Bealer K, Madden TL. 2009. BLAST+: architecture and applications. BMC Bioinformatics, 10:421.

Caravas J, Friedrich M. 2013. Shaking the Diptera tree of life: performance analysis of nuclear and mitochondrial sequence data partitions. Systematic Entomology, 38:93–103.

Cavener DR. 1987. Comparison of the consensus sequence flanking translational start sites in Drosophila and vertebrates. Nucleic Acids Research, 15:1353–1361.

Chamberlain JS, Gibbs RA, Ranier JE, Nguyen PN, Caskey CT. 1988. Deletion screening of the Duchenne muscular dystrophy locus via multiplex DNA amplification. Nucleic Acids Res, 16:11141–11156.

Chen F, Mackey AJ, Stoeckert CJ, Jr., Roos DS. 2006. OrthoMCL-DB: querying a comprehensive multi-species collection of ortholog groups. Nucleic Acids Res, 34:D363–368.

Chen F, Mackey AJ, Vermunt JK, Roos DS. 2007. Assessing performance of orthology detection strategies applied to eukaryotic genomes. PLoS One, 2:e383.

Chifman J, Kubatko L. 2014. Quartet inference from SNP data under the coalescent model. Bioinformatics, 30:3317–3324.

Chou J, Gupta A, Yaduvanshi S, Davidson R, Nute M, Mirarab S, Warnow T. 2015. A comparative study of SVDquartets and other coalescent-based species tree estimation methods. BMC Genomics, 16:S2.

Cock PJ, Antao T, Chang JT, Chapman BA, Cox CJ, Dalke A, Friedberg I, Hamelryck T, Kauff F, Wilczynski B, de Hoon MJ. 2009. Biopython: freely available Python tools for computational molecular biology and bioinformatics. Bioinformatics, 25:1422–1423.

DaCosta JM, Sorenson MD. 2016. ddRAD-seq phylogenetics based on nucleotide, indel, and presence-absence polymorphisms: Analyses of two avian genera with contrasting histories. Mol Phylogenet Evol, 94:122–135.

Dale R. 2013. gffutils. https://daler.github.io/gffutils/.

Doumith M, Day MJ, Hope R, Wain J, Woodford N. 2012. Improved multiplex PCR strategy for rapid assignment of the four major Escherichia coli phylogenetic groups. J Clin Microbiol, 50:3108–3110.

Drew RAI, Romig MC. 2013. Tropical Fruit Flies of South-East Asia (Tephritidae: Dacinae). Wallingford, CABI.

Drew RAI, Romig MC. 2016. Keys to the Tropical Fruit Flies of South-East Asia. Wallingford, CABI.

Drummond AJ, Suchard MA, Xie D, Rambaut A. 2012. Bayesian phylogenetics with BEAUti and the BEAST 1.7. Mol Biol Evol, 29:1969–1973.

Dupuis JR, Brunet BM, Bird HM, Lumley LM, Fagua G, Boyle B, Levesque R, Cusson M, Powell JA, Sperling FA. 2017. Genome-wide SNPs resolve phylogenetic relationships in the North American spruce budworm (Choristoneura fumiferana) species complex. Mol Phylogenet Evol, 111:158–168.

Edwards MC, Gibbs RA. 1994. Multiplex PCR: advantages, development, and applications. Genome Res, 3:S65–75.

Faircloth BC. 2017. Identifying conserved genomic elements and designing universal bait sets to enrich them. Methods in Ecology and Evolution, 8:1103–1112.

Faircloth BC, Branstetter MG, White ND, Brady SG. 2015. Target enrichment of ultraconserved elements from arthropods provides a genomic perspective on relationships among Hymenoptera. Mol Ecol Resour, 15:489–501.

Faircloth BC, Chang J, Alfaro ME. 2012a. TAPIR enables high-throughput estimation and comparison of phylogenetic informativeness using locus-specific substitution models. asXiv.

Faircloth BC, McCormack JE, Crawford NG, Harvey MG, Brumfield RT, Glenn TC. 2012b. Ultraconserved elements anchor thousands of genetic markers spanning multiple evolutionary timescales. Syst Biol, 61:717–726.

Faircloth BC, Sorenson L, Santini F, Alfaro ME. 2013. A Phylogenomic Perspective on the Radiation of Ray-Finned Fishes Based upon Targeted Sequencing of Ultraconserved Elements (UCEs). PLoS One, 8:e65923.

Fan JB, Chee MS, Gunderson KL. 2006. Highly parallel genomic assays. Nat Rev Genet, 7:632–644.

Geib SM, Calla B, Hall B, Hou S, Manoukis NC. 2014. Characterizing the developmental transcriptome of the oriental fruit fly, Bactrocera dorsalis (Diptera: Tephritidae) through comparative genomic analysis with Drosophila melanogaster utilizing modENCODE datasets. BMC Genomics, 15:942.

Gostel MR, Coy KA, Weeks A. 2015. Microfluidic PCR-based target enrichment: A case study in two rapid radiations of Commiphora (Burseraceae) from Madagascar. Journal of Systematics and Evolution, 53:411–431.

Grabherr MG, Haas BJ, Yassour M, Levin JZ, Thompson DA, Amit I, Adiconis X, Fan L, Raychowdhury R, Zeng Q, Chen Z, Mauceli E, Hacohen N, Gnirke A, Rhind N, di Palma F, Birren BW, Nusbaum C, Lindblad-Toh K, Friedman N, Regev A. 2011. Full-length transcriptome assembly from RNA-Seq data without a reference genome. Nat Biotechnol, 29:644–652.

Guindon S, Dufayard JF, Lefort V, Anisimova M, Hordijk W, Gascuel O. 2010. New algorithms and methods to estimate maximum-likelihood phylogenies: assessing the performance of PhyML 3.0. Syst Biol, 59:307–321.

Hahn C, Fromm B, Bachmann L. 2014. Comparative genomics of flatworms (Platyhelminthes) reveals shared genomic features of ecto- and endoparastic neodermata. Genome Biol Evol, 6:1105–1117.

Hamilton CA, Lemmon AR, Lemmon EM, Bond JE. 2016. Expanding anchored hybrid enrichment to resolve both deep and shallow relationships within the spider tree of life. BMC Evol Biol, 16:212.

Hedin M, Starrett J, Akhter S, Schonhofer AL, Shultz JW. 2012. Phylogenomic resolution of paleozoic divergences in harvestmen (Arachnida, Opiliones) via analysis of next-generation transcriptome data. PLoS One, 7:e42888.

Hedtke SM, Morgan MJ, Cannatella DC, Hillis DM. 2013. Targeted enrichment: maximizing orthologous gene comparisons across deep evolutionary time. PLoS One, 8:e67908.

Hendrichs J, Vera MT, De Meyer M, Clarke AR. 2015. Resolving cryptic species complexes of major tephritid pests. Zookeys:5-39.

Hosner PA, Faircloth BC, Glenn TC, Braun EL, Kimball RT. 2016. Avoiding Missing Data Biases in Phylogenomic Inference: An Empirical Study in the Landfowl (Aves: Galliformes). Mol Biol Evol, 33:1110–1125.

Johnson BR, Borowiec ML, Chiu JC, Lee EK, Atallah J, Ward PS. 2013. Phylogenomics resolves evolutionary relationships among ants, bees, and wasps. Curr Biol, 23:2058–2062.

Jones E, Oliphant E, Peterson P, al. e. 2001. SciPy: open source scientific tools for Python. http://www.scipy.org/.

Jones MR, Good JM. 2016. Targeted capture in evolutionary and ecological genomics. Mol Ecol, 25:185–202.

Katoh K, Standley DM. 2013. MAFFT multiple sequence alignment software version 7: improvements in performance and usability. Mol Biol Evol, 30:772–780.

Kawahara AY, Breinholt JW. 2014. Phylogenomics provides strong evidence for relationships of butterflies and moths. Proc Biol Sci, 281:20140970.

Keilwagen J, Wenk M, Erickson JL, Schattat MH, Grau J, Hartung F. 2016. Using intron position conservation for homology-based gene prediction. Nucleic Acids Res, 44:e89.

Kinde I, Wu J, Papadopoulos N, Kinzler KW, Vogelstein B. 2011. Detection and quantification of rare mutations with massively parallel sequencing. Proceedings of the National Academy of Sciences, 108:9530–9535.

Kou R, Lam H, Duan H, Ye L, Jongkam N, Chen W, Zhang S, Li S. 2016. Benefits and Challenges with Applying Unique Molecular Identifiers in Next Generation Sequencing to Detect Low Frequency Mutations. PLoS One, 11:e0146638.

Kristensen DM, Wolf YI, Mushegian AR, Koonin EV. 2011. Computational methods for Gene Orthology inference. Brief Bioinform, 12:379–391.

Krosch MN, Schutze MK, Armstrong KF, Graham GC, Yeates DK, Clarke AR. 2012. A molecular phylogeny for the Tribe Dacini (Diptera: Tephritidae): systematic and biogeographic implications. Mol Phylogenet Evol, 64:513–523.

Kumar S, Filipski AJ, Battistuzzi FU, Kosakovsky Pond SL, Tamura K. 2012. Statistics and truth in phylogenomics. Mol Biol Evol, 29:457–472.

Lanfear R, Frandsen PB, Wright AM, Senfeld T, Calcott B. 2017. PartitionFinder 2: New Methods for Selecting Partitioned Models of Evolution for Molecular and Morphological Phylogenetic Analyses. Mol Biol Evol, 34:772–773.

Larsson A. 2014. AliView: a fast and lightweight alignment viewer and editor for large datasets. Bioinformatics, 30:3276–3278.

Leache AD, Chavez AS, Jones LN, Grummer JA, Gottscho AD, Linkem CW. 2015. Phylogenomics of phrynosomatid lizards: conflicting signals from sequence capture versus restriction site associated DNA sequencing. Genome Biol Evol, 7:706–719.

Leblanc L, Hossain MA, Khan SA, Jose MS, Rubinoff D. 2013. A Preliminary Survey of the Fruit Flies (Diptera: Tephritidae: Dacinae) of Bangladesh. Proceedings of the Hawaiian Entomological Society, 45:51–58.

Leblanc L, San Jose M, Barr N, Rubinoff D. 2015. A phylogenetic assessment of the polyphyletic nature and intraspecific color polymorphism in the Bactrocera dorsalis complex (Diptera, Tephritidae). Zookeys:339–367.

Lemmon AR, Emme SA, Lemmon EM. 2012. Anchored hybrid enrichment for massively high-throughput phylogenomics. Syst Biol, 61:727–744.

Lemmon EM, Lemmon AR. 2013. High-Throughput Genomic Data in Systematics and Phylogenetics. Annual Review of Ecology, Evolution, and Systematics, 44:99–121.

Li L, Stoeckert CJ, Jr., Roos DS. 2003. OrthoMCL: identification of ortholog groups for Eukaryotic genomes. Genome Res, 13:2178–2189.

Magoc T, Salzberg SL. 2011. FLASH: fast length adjustment of short reads to improve genome assemblies. Bioinformatics, 27:2957–2963.

Markoulatos P, Siafakas N, Moncany M. 2002. Multiplex polymerase chain reaction: a practical approach. J Clin Lab Anal, 16:47–51.

Martin M. 2011. Cutadapt removes adapter sequences from high-throughput sequencing reads. EMBnet. J., 17:10–12.

McCormack JE, Faircloth BC, Crawford NG, Gowaty PA, Brumfield RT, Glenn TC. 2012. Ultraconserved elements are novel phylogenomic markers that resolve placental mammal phylogeny when combined with species-tree analysis. Genome Res, 22:746–754.

McCormack JE, Hird SM, Zellmer AJ, Carstens BC, Brumfield RT. 2013. Applications of next-generation sequencing to phylogeography and phylogenetics. Mol Phylogenet Evol, 66:526–538.

McCormack JE, Tsai WL, Faircloth BC. 2016. Sequence capture of ultraconserved elements from bird museum specimens. Mol Ecol Resour, 16:1189–1203.

McKinney W. 2010. Data Structures for Statistical Computing in Python. Proceedings of the 9th Python in Science Conference:51–56.

Minh BQ, Nguyen MA, von Haeseler A. 2013. Ultrafast approximation for phylogenetic bootstrap. Mol Biol Evol, 30:1188–1195.

Mirarab S, Warnow T. 2015. ASTRAL-II: coalescent-based species tree estimation with many hundreds of taxa and thousands of genes. Bioinformatics, 31:i44–52.

Moyle RG, Oliveros CH, Andersen MJ, Hosner PA, Benz BW, Manthey JD, Travers SL, Brown RM, Faircloth BC. 2016. Tectonic collision and uplift of Wallacea triggered the global songbird radiation. Nat Commun, 7:12709.

Nguyen LT, Schmidt HA, von Haeseler A, Minh BQ. 2015. IQ-TREE: a fast and effective stochastic algorithm for estimating maximum-likelihood phylogenies. Mol Biol Evol, 32:268–274.

Nylander JAA. 2016. catfasta2phyml. GitHub repository, https://github.com/nylander/catfasta2phyml.

O'Neill EM, Schwartz R, Bullock CT, Williams JS, Shaffer HB, Aguilar-Miguel X, Parra-Olea G, Weisrock DW. 2013. Parallel tagged amplicon sequencing reveals major lineages and phylogenetic structure in the North American tiger salamander (Ambystoma tigrinum) species complex. Mol Ecol, 22:111–129.

Papanicolaou A, Schetelig MF, Arensburger P, Atkinson PW, Benoit JB, Bourtzis K, Castanera P, Cavanaugh JP, Chao H, Childers C, Curril I, Dinh H, Doddapaneni H, Dolan A, Dugan S, Friedrich M, Gasperi G, Geib S, Georgakilas G, Gibbs RA, Giers SD, Gomulski LM, Gonzalez-Guzman M, Guillem-Amat A, Han Y, Hatzigeorgiou AG, Hernandez-Crespo P, Hughes DS, Jones JW, Karagkouni D, Koskinioti P, Lee SL, Malacrida AR, Manni M, Mathiopoulos K, Meccariello A, Murali SC, Murphy TD, Muzny DM, Oberhofer G, Ortego F, Paraskevopoulou MD, Poelchau M, Qu J, Reczko M, Robertson HM, Rosendale AJ, Rosselot AE, Saccone G, Salvemini M, Savini G, Schreiner P, Scolari F, Siciliano P, Sim SB, Tsiamis G, Urena E, Vlachos IS, Werren JH, Wimmer EA, Worley KC, Zacharopoulou A, Richards S, Handler AM. 2016. The whole genome sequence of the Mediterranean fruit fly, Ceratitis capitata (Wiedemann), reveals insights into the biology and adaptive evolution of a highly invasive pest species. Genome Biol, 17:192.

Paradis E, Claude J, Strimmer K. 2004. APE: Analyses of Phylogenetics and Evolution in R language. Bioinformatics, 20:289–290.

Peloso PLV, Frost DR, Richards SJ, Rodrigues MT, Donnellan S, Matsui M, Raxworthy CJ, Biju SD, Lemmon EM, Lemmon AR, Wheeler WC. 2016. The impact of anchored phylogenomics and taxon sampling on phylogenetic inference in narrow-mouthed frogs (Anura, Microhylidae). Cladistics, 32:113–140.

Philippe H, Brinkmann H, Lavrov DV, Littlewood DTJ, Manuel M, Worheide G, Baurain D. 2011. Resolving difficult phylogenetic questions: why more sequences are not enough. PLoS Biol, 9:e1000602.

Philippe H, Roure B. 2011. Difficult phylogenetic questions: more data, maybe; better methods, certainly. BMC Biology, 9:91.

Phuc HK, Ball AJ, Son L, Hanh NV, Tu ND, Lien NG, Verardi A, Townson H. 2003. Multiplex PCR assay for malaria vector Anopheles minimus and four related species in the Myzomyia Series from Southeast Asia. Medical and Veterinary Entomology, 17:423–428.

Pond SL, Frost SD, Muse SV. 2005. HyPhy: hypothesis testing using phylogenies. Bioinformatics, 21:676–679.

Portik DM, Smith LL, Bi K. 2016. An evaluation of transcriptome-based exon capture for frog phylogenomics across multiple scales of divergence (Class: Amphibia, Order: Anura). Mol Ecol Resour, 16:1069–1083.

Prum RO, Berv JS, Dornburg A, Field DJ, Townsend JP, Lemmon EM, Lemmon AR. 2015. A comprehensive phylogeny of birds (Aves) using targeted next-generation DNA sequencing. Nature, 526:569–573.

Python Software Foundation. 2017. Python Language Reference, version 3.6.

R Core Team. 2016. R: a language and environment for statistical computing. R Foundation for Statistical Computing, Vienna, Austria. URL https://www.r-project.org/.

Rambaut A, Drummond AJ. 2010. FigTree v1.4.2. Institute of Evolutionary Biology, University of Edinburgh. Available at: http://tree.bio.ed.ac.uk/software/figtree.

Rambaut A, Suchard MA, Xie D, Drummond AJ. 2014. Tracer v1.6, available from http://beast.bio.ed.ac.uk/Tracer.

Rezende VB, Congrains C, Lima AL, Campanini EB, Nakamura AM, Oliveira JL, Chahad-Ehlers S, Junior IS, Alves de Brito R. 2016. Head Transcriptomes of Two Closely Related Species of Fruit Flies of the Anastrepha fraterculus Group Reveals Divergent Genes in Species with Extensive Gene Flow. G3 (Bethesda), 6:3283–3295.

Richardson BA, Page JT, Bajgain P, Sanderson SC, Udall JA. 2012. Deep sequencing of amplicons reveals widespread intraspecific hybridization and multiple origins of polyploidy in big sagebrush (Artemisia tridentata; Asteraceae). Am J Bot, 99:1962–1975.

Rohland N, Reich D. 2012. Cost-effective, high-throughput DNA sequencing libraries for multiplexed target capture. Genome Res, 22:939–946.

Ruane S, Austin CC. 2017. Phylogenomics using formalin-fixed and 100+ year-old intractable natural history specimens. Mol Ecol Resour.

Sayyari E, Mirarab S. 2016. Fast Coalescent-Based Computation of Local Branch Support from Quartet Frequencies. Mol Biol Evol, 33:1654–1668.

Schrempf D, Minh BQ, De Maio N, von Haeseler A, Kosiol C. 2016. Reversible polymorphism-aware phylogenetic models and their application to tree inference. J Theor Biol, 407:362–370.

Schutze MK, Aketarawong N, Amornsak W, Armstrong KF, Augustinos AA, Barr N, Bo W, Bourtzis K, Boykin LM, CÁCeres C, Cameron SL, Chapman TA, Chinvinijkul S, ChomiČ A, De Meyer M, Drosopoulou E, Englezou A, Ekesi S, Gariou-Papalexiou A, Geib SM, Hailstones D, Hasanuzzaman M, Haymer D, Hee AKW, Hendrichs J, Jessup A, Ji Q, Khamis FM, Krosch MN, Leblanc LUC, Mahmood K, Malacrida AR, Mavragani-Tsipidou P, Mwatawala M, Nishida R, Ono H, Reyes J, Rubinoff D, San Jose M, Shelly TE, Srikachar S, Tan KH, Thanaphum S, Haq I, Vijaysegaran S, Wee SL, Yesmin F, Zacharopoulou A, Clarke AR. 2015. Synonymization of key pest species within the Bactrocera dorsalis species complex (Diptera: Tephritidae): taxonomic changes based on a review of 20 years of integrative morphological, molecular, cytogenetic, behavioural and chemoecological data. Systematic Entomology, 40:456–471.

Schutze MK, Bourtzis K, Cameron SL, Clarke AR, De Meyer M, Hee AKW, Hendrichs J, Krosch MN, Mwatawala M. 2017. Integrative taxonomy versus taxonomic authority without peer review: the case of the oriental fruit fly, Bactrocera dorsalis (Tephritidae). Systematic Entomology.

Schutze MK, Virgilio M, Norrbom A, Clarke AR. 2016. Tephritid Integrative Taxonomy: Where We Are Now, with a Focus on the Resolution of Three Tropical Fruit Fly Species Complexes. Annu Rev Entomol.

Segura MD, Callejas C, Fernández MP, Ochando MD. 2007. New contributions towards the understanding of the phylogenetic relationships among economically important fruit flies (Diptera: Tephritidae). Bulletin of Entomological Research, 96:279–288.

Sim SB, Calla B, Hall B, DeRego T, Geib SM. 2015. Reconstructing a comprehensive transcriptome assembly of a white-pupal translocated strain of the pest fruit fly Bactrocera cucurbitae. Gigascience, 4:14.

Sim SB, Geib SM. 2017. A Chromosome-Scale Assembly of the Bactrocera cucurbitae Genome Provides Insight to the Genetic Basis of white pupae. G3 (Bethesda), 7:1927–1940.

Stamatakis A. 2014. RAxML version 8: a tool for phylogenetic analysis and post-analysis of large phylogenies. Bioinformatics, 30:1312–1313.

Stiller M, Knapp M, Stenzel U, Hofreiter M, Meyer M. 2009. Direct multiplex sequencing (DMPS)--a novel method for targeted high-throughput sequencing of ancient and highly degraded DNA. Genome Res, 19:1843–1848.

Swofford DL. 2017. PAUP test-version. Available at https://people.sc.fsu.edu/~dswofford/paup_test/.

Tange O. 2011. GNU Parallel - the command-line power tool. The USENIX Magazine, February:42–47.

The Inkscape Team. 2017. Inkscape v0.91. Available at: https://inkscape.org/.

Townsend JP. 2007. Profiling phylogenetic informativeness. Syst Biol, 56:222–231.

Turner EH, Ng SB, Nickerson DA, Shendure J. 2009. Methods for genomic partitioning. Annu Rev Genomics Hum Genet, 10:263–284.

Uribe-Convers S, Settles ML, Tank DC. 2016. A Phylogenomic Approach Based on PCR Target Enrichment and High Throughput Sequencing: Resolving the Diversity within the South American Species of Bartsia L. (Orobanchaceae). PLoS One, 11:e0148203.

van der Walt S, Colbert SC, Varoquaux G. 2011. The NumPy array: a structure for efficient numerical computation. Computing in Science and Engineering, 13:22–30.

Vargas RI, Pinero JC, Leblanc L. 2015. An Overview of Pest Species of Bactrocera Fruit Flies (Diptera: Tephritidae) and the Integration of Biopesticides with Other Biological Approaches for Their Management with a Focus on the Pacific Region. Insects, 6:297–318.

Virgilio M, Jordaens K, Verwimp C, White IM, De Meyer M. 2015. Higher phylogeny of frugivorous flies (Diptera, Tephritidae, Dacini): localised partition conflicts and a novel generic classification. Mol Phylogenet Evol, 85:171–179.

White IM, Elson-Harris MM. 1992. Fruit Flies of Economic Significance: Their Indentification and Bionomics. Wallingford, UK, CABI International.

Wielstra B, Duijm E, Lagler P, Lammers Y, Meilink WR, Ziermann JM, Arntzen JW. 2014. Parallel tagged amplicon sequencing of transcriptome-based genetic markers for Triturus newts with the Ion Torrent next-generation sequencing platform. Mol Ecol Resour, 14:1080–1089.

Wiens JJ, Tiu J. 2012. Highly incomplete taxa can rescue phylogenetic analyses from the negative impacts of limited taxon sampling. PLoS One, 7:e42925.

Xi Z, Liu L, Davis CC. 2015. Genes with minimal phylogenetic information are problematic for coalescent analyses when gene tree estimation is biased. Mol Phylogenet Evol, 92:63–71.

Yourstone SM, Lundberg DS, Dangl JL, Jones CD. 2014. MT-Toolbox: improved amplicon sequencing using molecule tags. BMC Bioinformatics, 15:284.

